# A Machine Learning Approach Reveals CRISPR-Cas I-F as a Genomic Marker of Antibiotic Susceptibility in Uropathogenic *E. coli*

**DOI:** 10.1101/2025.10.13.681552

**Authors:** Alexandra M. Young, Peter Humburg, Fang Liu, Michael C. Wehrhahn, Alfred Tay, Stephen M. Riordan, Li Zhang

## Abstract

**Background:** Antimicrobial resistance (AMR) in *Escherichia coli* is a critical global health challenge, particularly in urinary tract infections, where first-line treatments are increasingly compromised. While horizontal gene transfer (HGT) via mobile genetic elements is a major driver of AMR, the genomic factors that may constrain resistance gene acquisition remain underexplored. CRISPR-Cas systems, which provide adaptive immunity against foreign DNA, could influence AMR dynamics, but their role in *E. coli* remains incompletely understood.

**Methods:** We conducted a comprehensive whole-genome analysis of uropathogenic *E. coli* isolates, including a newly sequenced collection from Australian clinical samples and an independent, globally sourced validation cohort. Antimicrobial susceptibility profiles were integrated with CRISPR-Cas subtype classification, resistance gene burden, and mobile element content. Elastic net regression, adaptive lasso, and tree-based machine learning models were used to identify genomic predictors of resistance, with performance validated across both datasets.

**Results:** CRISPR-Cas subtype I-F was consistently associated with susceptibility to antibiotics commonly acquired through HGT, including trimethoprim and ampicillin, and linked to lower ARG and MGE burden. In contrast, Type I-E arrays, especially when co-occurring with orphan I-F arrays, were associated with increased resistance. These associations remained robust after adjusting for phylogroup, plasmid content, and genomic background, and were validated across datasets.

**Conclusions:** Subtype-specific CRISPR-Cas systems shape antibiotic resistance profiles in *E. coli*, with Type I-F functioning as a potential genomic barrier to ARG acquisition. These findings highlight CRISPR array typing as a novel biomarker for AMR risk prediction and surveillance, and suggest new opportunities for leveraging CRISPR-based mechanisms to limit resistance propagation in clinical contexts.

## Background

The rise of antimicrobial resistance (AMR) is a global health crisis demanding innovative solutions, with recent projections estimating 8.22 million annual deaths by 2050 if left unaddressed ^1^. Urinary tract infections (UTIs) are among the most common bacterial infections worldwide, with over 400 million cases annually ^2^. *Escherichia coli* is the predominant causative agent, responsible for 65-75% of all cases ^3^. The increasing prevalence of antibiotic resistant *E. coli* strains, particularly those carrying multidrug-resistant (MDR) plasmids, underscores the urgency of addressing the genomic mechanisms underpinning resistance ^4–6^.

A key driver of AMR is horizontal gene transfer (HGT) and mobile genetic elements (MGEs). MGEs, including plasmids, insertion sequences, transposons and integrons, facilitate the capture, accumulation, and dissemination of resistance genes by promoting intracellular and intercellular DNA mobility ^7^. Plasmids are the primary vehicle for transferring ARGs between bacteria, including distantly related species ^8^. The widespread acquisition of MDR plasmids within pathogenic species has been cause for concern ^9^, with plasmid-mediated resistance to the last resort antibiotic colistin reported almost a decade ago ^10^. These mobile elements often carry multiple resistance genes as cassettes, contributing to the rapid spread of MDR.

While MGEs can occasionally provide adaptive benefits, most notably under antibiotic pressure, their accumulation poses a substantial evolutionary trade-off for bacterial hosts. Despite being non-living, MGEs behave as selfish genetic elements, with evolutionary trajectories distinct from, and often at odds with, those of their host ^11^. Crucially, conjugative plasmid acquisition itself frequently incurs a measurable short-term fitness cost regardless of the genetic cargo^12^. Beyond this intrinsic cost, plasmid replication imposes a metabolic burden^13^, and foreign DNA can disrupt regulatory balance^14^, interfere with resource allocation^15^, or instigate antagonism between co-resident MGEs^16^. These risks of genomic instability and broader metabolic consequences might be outweighed in an environment with the intense selective pressure of antibiotic exposure, where the loss of host defences may enable gene flow necessary for survival ^17,18^.

The ongoing exchange of genetic material via MGEs is therefore a critical driver of the development and spread of AMR among bacteria. While mechanisms of resistance acquisition are well characterised, comparatively little is known about the genomic factors that may act to limit the uptake or retention of resistance determinants, particularly under natural selective pressures. Identifying such limiting factors, which may serve as barriers to resistance dissemination, is essential to understanding how AMR spread may be restrained. The archaeal and bacterial adaptive immune system CRISPR represents perhaps the most important genetic defence to MGE acquisition. These CRISPR systems use short RNA molecules, derived from previous encounters with foreign genetic material, to guide Cas proteins in recognising and disabling invading MGEs such as bacteriophages and plasmids (reviewed in ^19^). In *E. coli* CRISPR arrays are most commonly classified as Type I-E or I-F ^20^, although strains have been identified with orphan I-F arrays lacking nearby *cas* genes which are thought to be nonfunctional ^21^.

Bacterial CRISPR-Cas systems are hypothesised to limit the spread of AMR genes by preventing the horizontal transfer of mobile resistance elements. However, this protective effect is not consistently observed. While it may be expected that strains with active CRISPR arrays would carry fewer plasmids, this correlation is not always evident ^22^. Touchon et al. ^23^ reported no association between CRISPR and plasmid, integron or ARG presence in *E. coli*, while others have noted differences in antimicrobial susceptibility based on CRISPR type ^24^. This suggests a complex interplay between CRISPR array subtype, MGEs, and AMR.

To clarify these conflicting findings and assess their global relevance, we performed a genome-wide analysis of *Escherichia coli* isolates from urinary tract infections. Using a newly sequenced dataset of Australian clinical isolates, we integrated CRISPR-Cas subtyping, antimicrobial susceptibility, resistance gene burden, and mobile element profiling. We then validated these associations in an independent, globally sourced dataset. By combining whole-genome sequencing with statistical modelling and interpretable machine learning, our study investigates how specific CRISPR-Cas subtypes influence resistance phenotypes and evaluates their potential as genomic markers for AMR risk stratification.

## Methods

### Clinical isolates and phenotypic resistance profiling

Clinical *E. coli* isolates were obtained from urine samples processed by Douglass Hanly Moir Pathology in 2023 during routine diagnostic procedures. To ensure a balanced representation of resistance phenotypes, isolates were selected to include approximately equal numbers of strains susceptible and resistant to trimethoprim. Selection was performed without prior knowledge of other genomic or clinical features to minimise selection bias.

Antimicrobial susceptibility was determined using the calibrated dichotomous sensitivities disc diffusion method and minimum inhibitory concentrations (MICs) were interpreted according to European Committee on Antimicrobial Susceptibility Testing (EUCAST) guidelines (v13.0). Isolates were tested for sensitivity to trimethoprim, ampicillin, amoxicillin-clavulanate, cefalexin, norfloxacin, nitrofurantoin and extended spectrum β-lactamase (ESBL) production.

### Whole genome sequencing and assembly

#### Short-read Illumina sequencing and assembly

Genomic DNA from the clinical *E. coli* isolates described above was extracted and sequenced as previously described ^25^. Briefly, DNA was extracted using the Gentra Puregene Yeast/Bacteria Kit (Qiagen, Australia) according to manufacturer’s instructions. DNA libraries were sequenced using the Illumina MiSeq Benchtop Sequencer at the Marshall Centre for Infectious Diseases Research and Training, University of Western Australia.

Raw reads were quality-trimmed with fastp (v0.12.4) ^26^ using default settings, and de novo assembly was performed using Shovill (v1.1.0) (https://github.com/tseemann/shovill) applying a minimum contig length of 500 bp and a minimum coverage threshold of 5.0. Assembly quality was assessed using QUAST (v5.2.0) ^27^. Assembly statistics for all short-read genomes are available in Supplementary Table 1.

Species confirmation was conducted via ribosomal multilocus sequence typing (rMLST) using the PubMLST database ^28,29^, and genome annotation was performed with PROKKA (v1.14.6) ^30^.

#### Long-read Nanopore sequencing and hybrid assembly

To further characterise the genomic context of resistance genes, long-read sequencing was performed on two representative isolates using Oxford Nanopore Technology. These isolates were selected to reflect distinct resistance profiles, with one resistant to trimethoprim alone and another with a multidrug-resistant phenotype including ampicillin, amoxicillin– clavulanate, and cefalexin.

Genomic DNA was extracted by phenol–chloroform purification, quantified, and prepared with the Q20+ ligation sequencing kit (SQK-LSK114, motor E8.2). Libraries were loaded onto R10.4.1 flow-cells (FLO-MIN114) and sequenced on an Oxford Nanopore MinION Mk1B at the Ramaciotti Centre for Genomics, University of New South Wales. Real-time base-calling was carried out in high-accuracy (HAC) mode with Guppy v6.5.7 using the 400 bp s□^1^ R10.4.1/E8.2 model. Raw read metrics were determined with NanoStat (v1.6.0) ^31^.

Sequenced raw reads were quality-filtered with Filtlong (v0.2.1) (github.com/rrwick/Filtlong), discarding reads less than 1 kb and the lowest-quality 5%. Filtered reads were subsampled into 12 independent subsets to generate candidate assemblies for Trycycler (v0.5.4) ^32^, which generates a consensus sequence from multiple assemblies. Candidate assemblies were generated using one of Flye (v2.9.2) ^33^, Raven (v1.8.3) ^34^ or Miniasm/Minipolish (v0.3/v0.1.3) ^35^. The Trycycler consensus contigs were polished with Medaka (v1.2.2) (https://github.com/nanoporetech/medaka) utilising the r1041_e82_400bps_hac_v4.2.0 model, followed by a second polishing step using trimmed Illumina reads and Polypolish (v0.5.0) ^36^. Final assemblies were visualised using the Proksee web tool ^37^. Sequencing and assembly metrics for the hybrid genomes of R52 and R60 are provided in Supplementary Table 2.

### Phylogenetic analysis

Core-genome alignments were generated using Roary (v3.13.0) ^38^ and approximate maximum-likelihood phylogenetic trees were constructed using FastTree (v2.1.11) ^39^ as previously described ^40^. Trees were visualised using the R packages (via RStudio v1.4.1717) ggtree (v3.2.1) ^41^ and ComplexHeatmap (v2.15.4) ^42^ .

### Gene screening

The *E. coli* phylogroups were determined *in silico* using the ClermonTyping web tool (v23.06) ^43^. Sequences types (STs) were determined using PubMLST ^29^ using the Achtman scheme, with the Pasteur scheme used for isolates which did not yield an ST. ARGs were identified with RGI (v6.0.2) referencing CARD ^44^. The presence of any CRISPR-Cas systems were detected and typed using CRISPRCasTyper (v1.8.0) ^45^. Mobile elements were identified using MobileElementFinder (v1.1.2) ^46^ IntegronFinder (v2.0.2) ^47^ and Plasmer (v23.04.20) ^48^.

### Global isolate validation

A total of 426 Illumina paired-end raw reads from urinary *E. coli* were obtained from the Sequence Read Archive (SRA) database. All reads had corresponding antimicrobial susceptibility results and were obtained from human urine samples.

Genome assembly and quality checks were performed as for the isolates sequenced within this study. Strains were filtered using a combination of rMLST and the criteria for *E. coli* used by Enterobase requiring a total length between 3.7 – 6.4Mbp, N50 of ≥20□kb and ≤800 contigs ^49^. Following filtering there were 303 isolates obtained from 7 countries, with the majority originating from the United States (n=194, 64.0%), Australia (n=67, 22.1%) and the United Kingdom (n=27, 8.9%). Isolate accession numbers and assembly quality are available in Supplementary Table 3 and the country and year of isolation, as well as the antimicrobial susceptibility testing results are available in Supplementary Table 4.

### Regularised regression: CRISPR array types and antimicrobial resistance prediction

We used elastic net regression followed by adaptive lasso to identify and select features associated with phenotypic antimicrobial resistance in uropathogenic *E. coli*. Elastic net regression was selected due to the high correlation present between genetic predictors; this method is effective in handling multicollinearity by shrinking coefficients of correlated variables. Adaptive lasso was then applied to enhance model interpretability. The models were trained using the data generated within this study, while the global dataset of human urine *E. coli* isolates was used for independent validation.

Predictors included within the model included phylogroups, plasmid types, CRISPR array types and specific ARGs. Predictors were filtered prior to analysis to only include those which are present in more than 5% of isolates within each dataset. Two continuous variables, ARG and MGE counts, were converted to binary by setting a cutoff based on the distribution of the data. For MGEs, a value of ≥21 was chosen, which was above the mean for both independent datasets, and for ARGs, a cutoff of ≥56 was selected, reflecting a clear division in the distribution between isolates. Due to the limited number of isolates exhibiting phenotypic resistance to amoxicillin-clavulanate, cefalexin, and norfloxacin, only trimethoprim and ampicillin resistance were included in the regression models. This approach was necessary to ensure sufficient statistical power for identifying reliable resistance-associated features.

Due to the variation in national prescribing practices globally ^50,51^, reported phenotypic resistance to trimethoprim-sulfamethoxazole was used for the global dataset, while the data generated in this study reported resistance to trimethoprim alone.

Given our focus on the interaction between CRISPR array types and AMR, we removed regularisation penalties for these predictors to ensure they were fully represented in all modelling.

Elastic net regression was applied with equal weighting between ridge and lasso regularisation using the glmnet package (v4.1-7) ^52^. The regularisation strength was optimised using 8-fold cross-validation to minimise cross-validation error based on the area under the receiver operating characteristic (ROC) curve (AUC). The non-zero coefficients identified by elastic net were retained, and their magnitudes were used to derive weights for the adaptive lasso penalty. Adaptive lasso was also implemented using glmnet ^52^, with hyperparameters optimised through another round of 8-fold cross-validation to minimise the cross-validation error based on AUC.

To evaluate the predictive performance of the model on the independent global validation dataset, we applied Youden’s J statistic to determine the optimal classification threshold. This threshold maximises the sum of sensitivity and specificity, providing a balance between correctly identifying resistant and susceptible isolates.

Model performance for the training data was evaluated using AUC. For the independent global validation dataset, performance was assessed using AUC, balanced accuracy, specificity, sensitivity, F1 score, and confusion matrices. The F1 score, defined as the harmonic mean of precision and sensitivity, was used to assess the balance between the model’s ability to identify true positives and avoid false positive.

### Extreme gradient boosting: genomic features and antimicrobial resistance prediction

Extreme gradient boosting (XGBoost) was conducted using the xgboost (v1.6.0.1) ^53^ and caret (v6.0-94) ^54^ R packages to assess the overall predictive capacity of genomic features for antimicrobial resistance. We employed 8-fold cross-validation and AUC as the primary performance metric. Key hyperparameters, including learning rate, tree depth, number of boosting rounds, and regularisation terms were optimised using a grid search approach to improve model performance. The full hyperparameter grid is provided in Supplementary Table 5. The predictors, filtering and training dataset generated within this study were identical to those used for the regularised regression models described above.

To evaluate the generalisability of the XGBoost model, the independent, geographically diverse dataset of human urine *E. coli* isolates was used for external validation. For the global predictions, the probability outputs from XGBoost were reclassified using the optimal threshold derived from Youden’s J statistic. Model performance for the training data was evaluated using AUC. Performance on the global validation dataset was assessed using AUC, balanced accuracy, specificity, sensitivity, F1 score, and confusion matrices.

### Decision tree modelling: interpretable prediction of antimicrobial resistance

Decision tree modelling was performed using the rpart (v4.1.21) (https://github.com/bethatkinson/rpart) and caret (v6.0-94) ^54^ packages to assess the predictive performance of genomic features for antimicrobial resistance. The same predictors, filtering criteria, and training dataset described for the regularised regression and XGBoost models were used here to ensure consistency across modelling approaches. Decision trees were selected as an interpretable, non-parametric alternative to more complex models, allowing for direct visualisation of feature-based splits associated with resistance outcomes. Model complexity was controlled by tuning the complexity parameter, which determines the minimum improvement in model fit required for a split to be retained. This was optimised using 8-fold cross-validation to maximise AUC, and the optimal value was selected based on cross-validated performance.

As with XGBoost, the final model was validated on the independent global dataset of *E. coli* isolates. Probabilistic outputs were converted to binary predictions using a classification threshold defined by Youden’s J statistic. Model performance was assessed using AUC on the training data, while external validation was evaluated using AUC, balanced accuracy, specificity, sensitivity, F1 score, and confusion matrices.

### Statistical analysis

Pairwise comparisons were assessed using Fisher’s exact test with correction for multiple comparisons using the Benjamini–Hochberg procedure. Gene count comparisons were evaluated using Dunn’s test and the Holm adjustment for multiple comparisons. All reported *p* values are adjusted for multiple comparisons, with statistical significance determined at α = 0.05. Statistical analyses were performed using R software (via RStudio v1.4.1717).

## Results

To investigate predictors of antimicrobial resistance in *E. coli*, we analysed two independent datasets comprising global genomes and a novel collection of 91 *E. coli* isolates sequenced from Australian predominately community-acquired urinary tract infections. These newly sequenced isolates were selected to ensure a balanced representation of trimethoprim susceptibility and served as a primary dataset for model training and feature discovery, with findings validated using the external global cohort.

### High prevalence of trimethoprim-ampicillin co-resistance observed in community-acquired UTI isolates

Following species confirmation and quality control of the newly sequenced Australian isolates, 49 (53.8%) were identified as trimethoprim-resistant and 42 (46.2%) as susceptible, forming the basis of downstream genomic analysis.

Of the 91 isolates, 51.6% were resistant to ampicillin (n=47), 6.6% (n=6) were resistant to amoxicillin-clavulanate, 7.7% (n=7) displayed resistance to cefalexin, 6.6% (n=6) to norfloxacin and 2.2% (n=2) were ESBL positive (Supplementary Fig. 1). There were 32 strains sensitive to all antibiotics tested, and no isolates displayed resistance to nitrofurantoin. Co-resistance between trimethoprim and ampicillin was common, 75.5% (n=37/49) of trimethoprim resistant isolates were also resistant to ampicillin, and 78.7% (n=37/47) of ampicillin resistant strains were also resistant to trimethoprim.

The majority of clinical samples were obtained in community settings (n=86, 94.5%), with three from aged care facilities (3.3%) and two from hospitals (2.2%) (Supplementary Table 6). The isolates were predominantly from females (n=75, 82.4%), with a mean patient age of 60 years. Most were over the age of 50 (n=64, 70.3%), and 59.3% (n=54) were over 60. Only two patients were under the age of 13 (2.2%).

### MDR plasmids harbor co-localised resistance genes within composite transposons

To investigate the genetic basis of the observed trimethoprim-ampicillin co-resistance, we examined the plasmid-associated resistance genes in our *E. coli* isolates. Resistance to trimethoprim was positively associated with 20 ARGs spanning multiple antibiotic classes (Fig. 1). These included genes conferring resistance to trimethoprim (*dfrA1, dfrA12, dfrA17*) and sulfonamides (*sul1, sul2*), which are commonly co-located on mobile resistance plasmids ^55,56^. Notably, genes conferring β-lactam resistance (*blaEC-8*, *blaTEM-1*), a fluoroquinolone resistance mutation (*parC*), and determinants against aminoglycosides, macrolides, tetracyclines, and fosfomycin were also enriched, reflecting a broad MDR profile associated with trimethoprim resistance.

**Figure 1.**
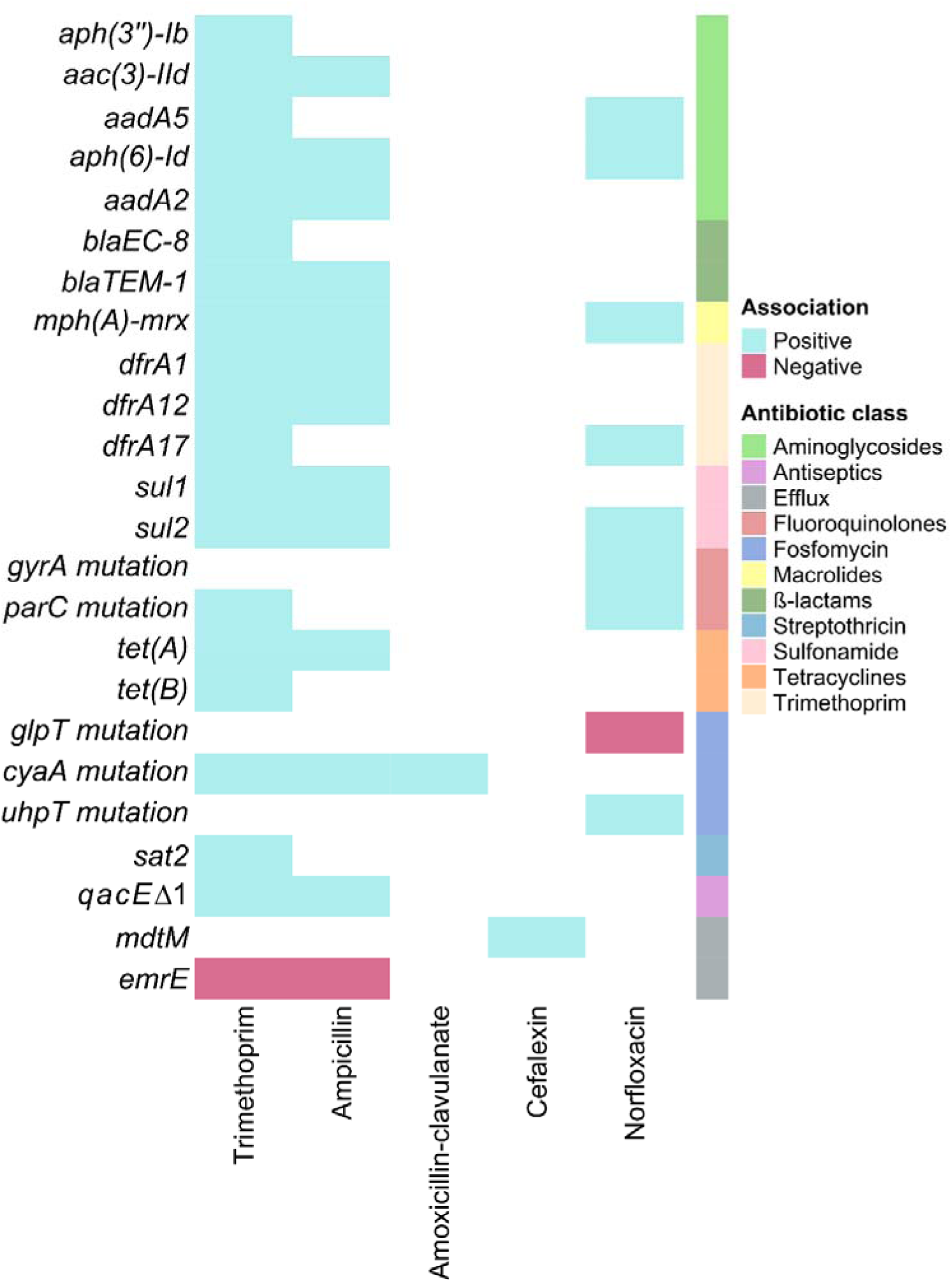
Statistically significant associations between phenotypic antibiotic resistance and resistance genes. Statistically significant associations between antibiotic resistance genes and phenotypic resistance to trimethoprim, ampicillin, amoxicillin-clavulanate, cefalexin, and norfloxacin were determined using Fisher’s exact test. P-values were adjusted for multiple comparisons using the Benjamini-Hochberg method. Positive associations indicate that the presence of the gene is correlated with resistance to the antibiotic and depicted in blue, while negative associations are shown in pink. All genes positively associated with ampicillin resistance were also linked to trimethoprim resistance, suggesting a strong genetic basis for co-resistance.

All 13 AMR genes positively associated with ampicillin resistance were also significantly associated with trimethoprim resistance (Fig. 1). Of these shared genes, only *cyaA* encoding adenylate cyclase, was chromosomally encoded; all others were plasmid-borne. These findings suggest that co-resistance is largely driven by shared carriage of resistance determinants on mobile elements.

To resolve the genomic architecture of these co-localised resistance genes, we performed long-read sequencing and hybrid assembly on two *E. coli* isolates, R52 and R60, selected to represent distinct resistance phenotypes (see Methods). R52 exhibited resistance to trimethoprim alone, while R60 displayed a broader MDR profile, including resistance to ampicillin, amoxicillin–clavulanate, and cefalexin.

Hybrid assembly revealed distinct conjugative MDR plasmids in each strain, as indicated by the presence of *tra* operons, suggesting potential for HGT. Strain R52 carried a 119kb FIA plasmid characterised by ARGs integrated within transposable elements, including a composite transposon carrying *tet(A), tet(R), sul1, qacE*Δ*1, ant(3”)-IIa* and *dfrA1,* as well as the enterotoxin protein TieB (*senB*) (Fig. 2a). The plasmid contained an additional composite transposon carrying *blaTEM-1*. This plasmid closely resembled a previously deposited unnamed 137 kb plasmid sequence (CP163781.1; 99.96% identity, 100% coverage) from *E. coli* strain OXEC-19. We designated the novel plasmid pR52-MDR1 for this study.

**Figure 2.**
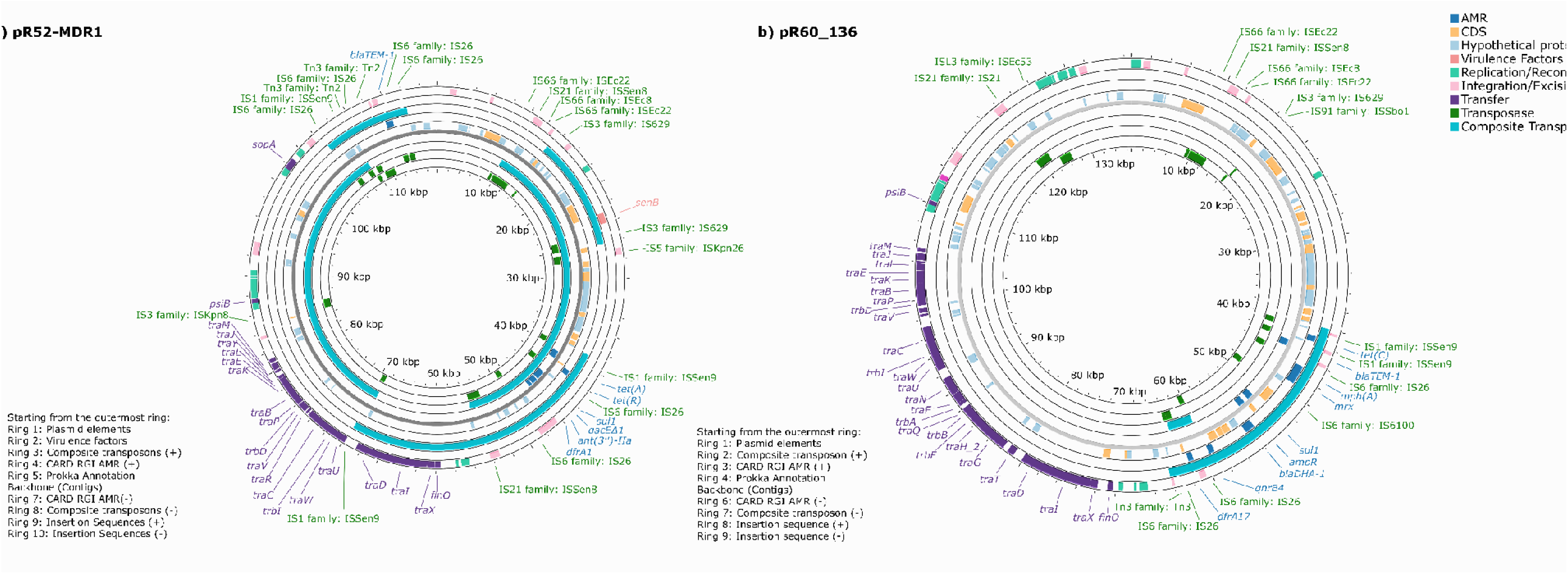
Genetic organisation of antimicrobial resistance genes on MDR plasmids identified by long-read sequencing. **a)** A 119kb FIA plasmid, pR52-MDR1, was identified in strain R52 carrying a variety of antimicrobial resistance genes. The host strain R52 did not possess any CRISPR arrays. ARGs are integrated within transposable elements. The presence of *sul1* and *dfrA1* aligns with the observed trimethoprim resistance in this strain. **b)** Strain R60 harboured a 136kb FII conjugative plasmid, pR60_136kb_FII. The host strain possessed type I-E and orphan I-F CRISPR arrays. All resistance genes are clustered within a hypermobile region of overlapping composite transposons, suggesting a mechanism for co-transfer. The presence of *blaTEM-1, blaDHA-1, dfrA17, sul1, and tet(C)* is consistent with the strain’s MDR profile, including resistance to trimethoprim, ampicillin, amoxicillin with clavulanate, and cefalexin.

Strain R60 harboured a 136kb FII plasmid, exhibiting high similarity to the 134kb pAVS0973-A (CP124472.1; 99.98% identity, 91% coverage. Notably in this plasmid, all resistance genes-including *dfrA17, qnrB4, blaDHA-1, ampR, sul1, mph(A)-mrx, blaTEM-1* and *tet(C)*, were co-located within a hypermobile region containing overlapping composite transposons (Fig. 2b), facilitating potential co-transfer of these genes.

Detailed structural maps of the composite transposon regions are provided in Supplementary Fig. 2.

### Phylogenetic distribution of antibiotic resistance and CRISPR systems

The core-genome tree saw isolates clustered by phylogroup (Fig. 3). The majority of isolates belonged to the B2 phylogroup (n=59, 64.8%), followed by D (n=16, 17.6%), B1 (n=6, 6.6%), A (n=5, 5.5%), F (n=4, 4.4%) and a single isolate in phylogroup C. We did not detect significant associations between phylogroup and phenotypic antibiotic resistance.

**Figure 3.**
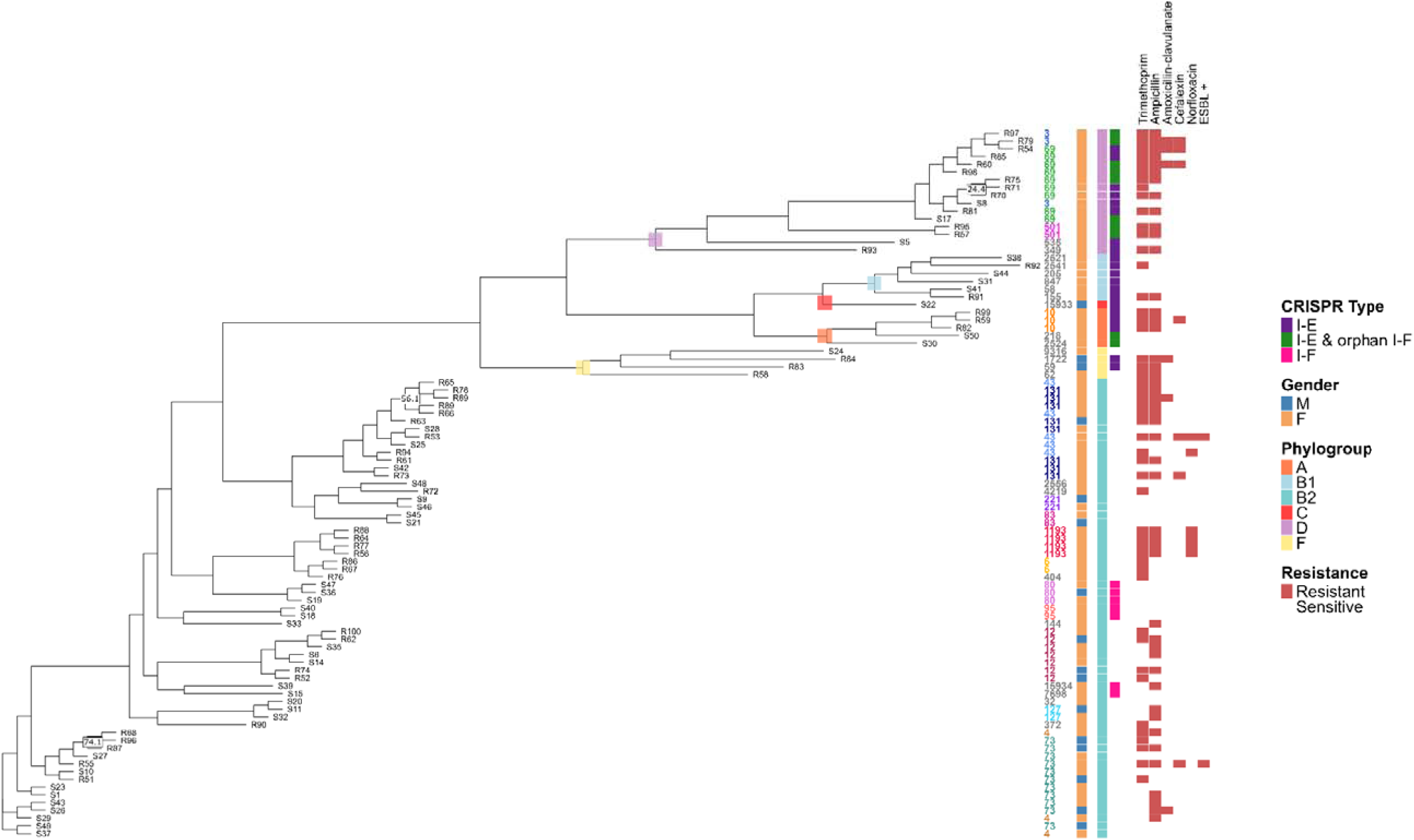
Core genome phylogeny of urinary pathogenic *E.coli* isolates reveals associations between phylogroup, sequence type, CRISPR array type, and phenotypic resistance. A core genome phylogeny was generated using Roary and FastTree. Isolates are clustered by phylogroup, which is indicated both on the branching nodes and the adjacent heatmap. Only bootstrap values less than 80 are displayed, all others are above this cutoff. The following characteristics are displayed for each isolate adjacent to the tree (from left to right): strain sequence type (ST), patient gender, phylogroup, CRISPR array type, and resistance phenotypes. See the accompanying key for a complete description of the colours representing each trait. Notably, the majority of strains belong to phylogroup B2 (n=59, 64.8%) and the type I-F CRISPR arrays were confined to this group, with B2 isolates either possessing the I-F array or no array at all.

A total of 37 different STs were identified, with 23 represented by a single isolate. The most frequent were ST73 (n=11, 12.1%), ST69 (n=9, 9.9%), ST131 (8, 8.8%), ST12 (n=7, 7.7%), ST43 (n=5, 5.5%) and ST1193 (n=4, 4.4%), which collectively accounted for 48.4% of all isolates. Notably, ST1193 strains consistently displayed resistance to trimethoprim, ampicillin and norfloxacin, with a significant association noted between ST1193 and norfloxacin resistance (p < 0.001). This may reflect the acquisition and maintenance of resistance elements and MDR plasmids, as well as the universal fluoroquinolone resistance present within the ST1193 lineage ^57^.

The presence of CRISPR systems also varied among *E. coli* strains (Fig. 3). The majority of strains lacked CRISPR arrays entirely (*n*=54, 59.3%), while others possessed complete type I-E arrays (n=20, 22.0%), complete I-F arrays (n=7, 7.7%), or a combination of complete I-E and orphan I-F arrays (n=10, 11.0%).

In both the local and global datasets, orphan I-F arrays were almost exclusively found in isolates that also carried complete I-E systems. In this study, we refer to this co-occurrence as a distinct CRISPR subtype.

This pattern reflects the phylogenetic structuring of CRISPR systems in *E. coli*, which has been observed in prior studies ^58^. CRISPR type I-E systems were negatively associated with the B2 phylogroup (*p* < 0.001), and positively associated with phylogroups A (*p* = 0.013), B1 (*p* = 0.001) and D (*p* = 0.037). Conversely, the absence of CRISPR arrays was positively associated with phylogroup B2 (*p* < 0.001), and negatively associated with groups A (*p* = 0.003), B1 (*p* < 0.02) and D (*p* < 0.001). Complete I-E arrays with orphan I-F were more common in phylogroup D (*p* < 0.001). All complete type I-F arrays were restricted to phylogroup B2, although this association was not statistically significant (*p* = 0.089).

### Complete Type I-F CRISPR system: a marker of reduced resistance

Clear associations were observed between the possession of a complete I-F system and AMR (Fig. 4). Strains with type I-F arrays harboured significantly fewer ARGs compared to those without CRISPR systems (p = 0.009), further suggesting a potential role in limiting the acquisition or retention of resistance genes. Additionally, the presence of a type I-F array was positively associated with trimethoprim susceptibility (p = 0.003).

**Figure 4.**
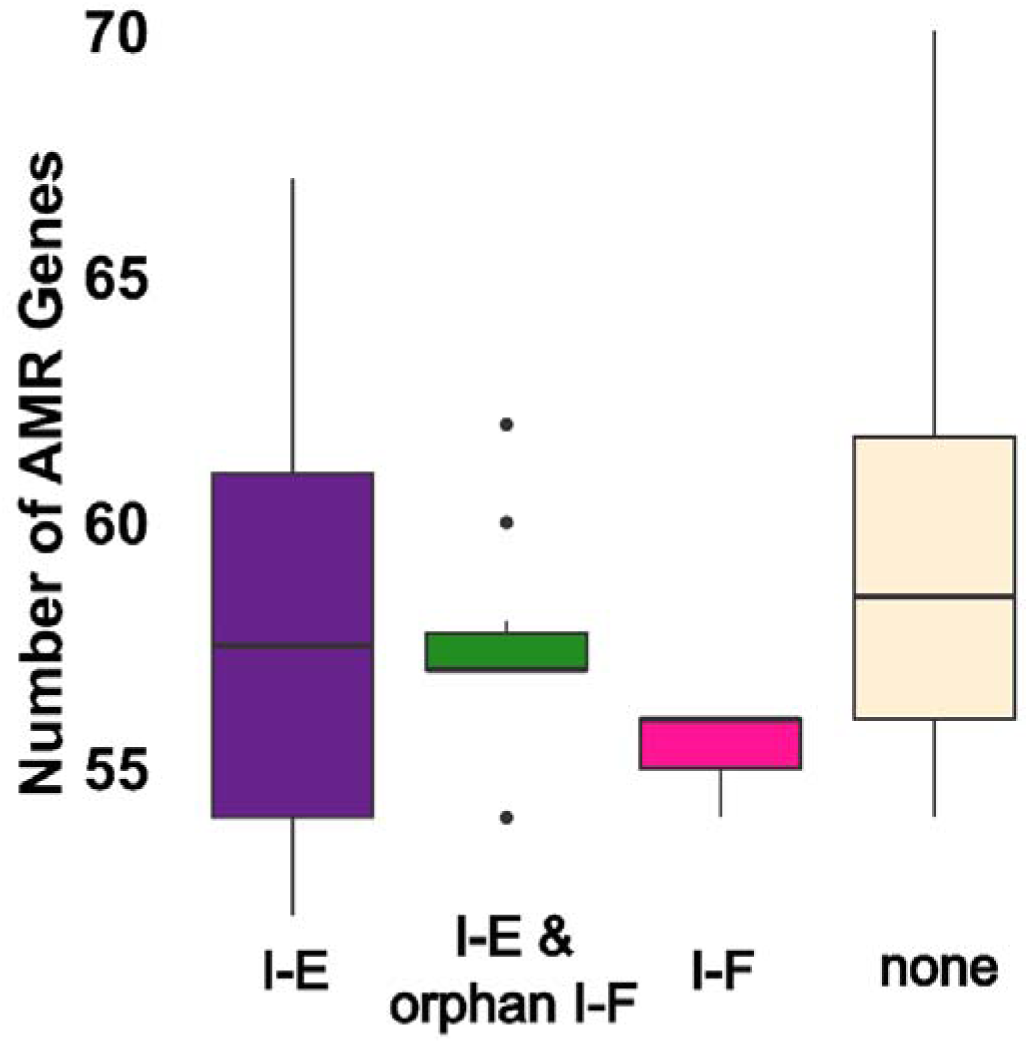
Distribution of antibiotic resistance genes across different CRISPR array types. The boxplots display the median number of antibiotic resistance genes and the interquartile range for each CRISPR array type. Strains harbouring a type I-F CRISPR array possessed significantly fewer AMR genes than strains lacking a CRISPR array (Dunn’s test with Holm adjustment for multiple comparisons, p=0.009).

Ampicillin resistance was notably rare among isolates with a type I-F system, contrasting with its general prevalence of >50% within this dataset. Among the type I-F strains, only one isolate exhibited resistance to ampicillin, while all others were pan-susceptible to the antibiotics tested. This observation was investigated further in the context of *blaTEM-1* prevalence. The *blaTEM-1* gene, encoding a common β-lactamase responsible for ampicillin resistance, was strikingly absent in strains possessing a type I-F CRISPR system. The *blaTEM-1* gene is often found on mobile genetic elements such as plasmids and transposons, facilitating its horizontal transfer between bacteria ^59^. Within the local dataset, 33% (n=30) of strains contained the gene. Among the 30 *blaTEM-1* positive strains, 10% (n=3) had complete I-E and orphan I-F arrays, 23.3% (n=7) had the I-E type and the remaining 66.7% (n=20) did not have a CRISPR system.

### Global validation confirms Type I-F CRISPR systems are linked to antimicrobial susceptibility and fewer mobile elements

Analysis of global *E. coli* strains isolated from urine revealed a similar CRISPR distribution pattern to the Australian dataset. Most global strains lacked CRISPR arrays (n=161, 53.5%), followed by those with type I-E (n=58, 19.3%), type I-F (n=50, 16.6%), and I-E arrays with orphan I-F (n=16, 5.3%) (Fig. 5). As in the local dataset, the majority of isolates belonged to phylogroup B2 (n=215, 71.4%), with smaller proportions in groups D (n=35, 11.6%), A (n=21, 7.0%), and B1 (n=20, 6.6%).

**Figure 5.**
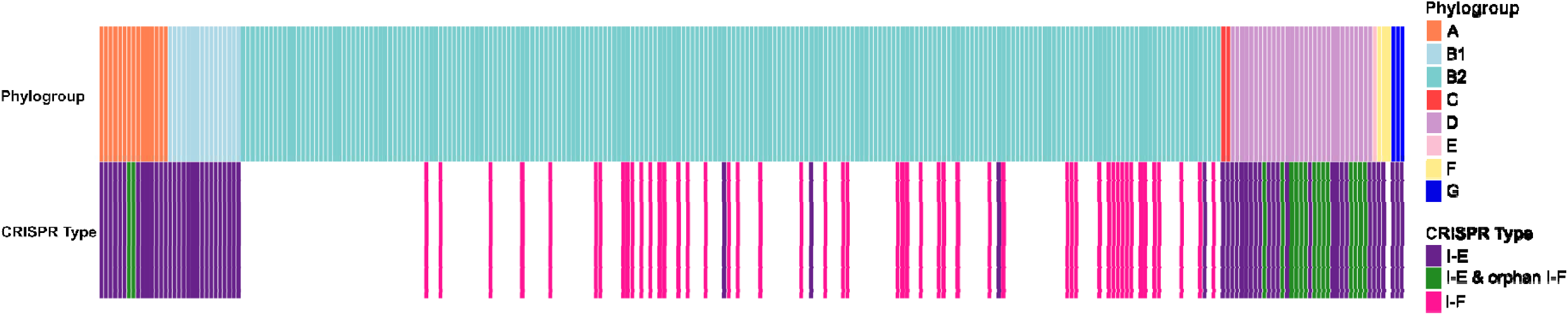
Distribution of phylogroups and CRISPR array types in the global uropathogenic *E. coli* isolate dataset. Each column represents a single isolate, with phylogroups represented in the top row and CRISPR array type in the bottom row. See the key for a complete description of the colours representing each phylogroup and CRISPR array type.

The global dataset supported the observed phylogenetic associations with CRISPR types. Type I-F CRISPR systems were exclusively found in phylogroup B2 (p < 0.001), while the absence of CRISPR arrays was also positively associated with B2 (p < 0.001) and negatively associated with groups A and B1 (p < 0.001). Type I-E arrays were more common in phylogroups A, B1, D (all p < 0.001), and G (p = 0.01), and I-E with orphan I-F arrays were strongly associated with phylogroup D (p < 0.001).

The association between type I-F CRISPR arrays and reduced antibiotic resistance was further validated. Strains possessing complete I-F systems harboured significantly fewer ARGs than those with I-E systems, I-E and orphan I-F, or no CRISPR arrays (p< 0.001) (Fig. 6a). Similarly, type I-F strains contained fewer MGEs compared to all other categories (p < 0.001) (Fig. 6b).

**Figure 6.**
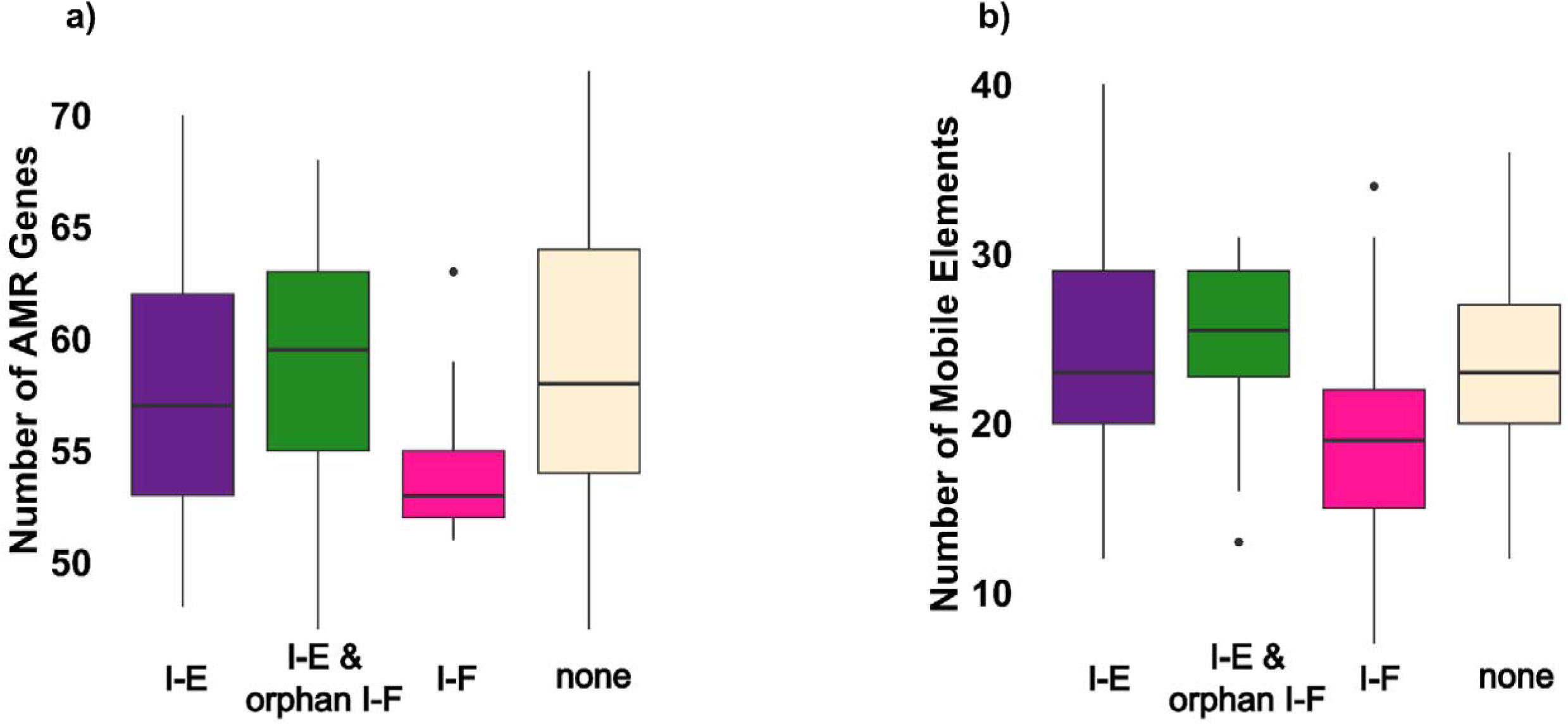
Impact of CRISPR-Cas systems on the acquisition of antibiotic resistance and mobile genetic elements in global *E. coli* isolates. Boxplots show distributions across CRISPR array type. Statistical comparisons were performed using Dunn’s test with Holm adjustment for multiple comparisons. **a)** Strains possessing complete I-F systems harboured significantly fewer ARGs than those with I-E systems, I-E and orphan I-F, or no CRISPR arrays (all p < 0.001. **b)** Type I-F strains contained fewer MGEs compared to all other categories (all p < 0.001).

Resistance phenotypes also varied by CRISPR type. The presence of a type I-F CRISPR array was negatively associated with resistance to multiple antibiotics, including ampicillin, cefotaxime, ciprofloxacin, gentamicin, and trimethoprim with sulfamethoxazole. Conversely, the absence of CRISPR arrays was associated with resistance against ampicillin, amoxicillin-clavulanate, gentamicin, and norfloxacin.

Further analysis of the strains encoding *blaTEM-1* reinforced the association observed within the local dataset. Among the 135 *blaTEM-1* positive isolates, only 7.4% (n=10) had type I-F CRISPR systems. The majority of *blaTEM-1* isolates did not possess a CRISPR system at all (63.0%, n=85), while 18.5% (n=25) had I-E systems and 8.1% had I-E and orphan I-F arrays. This inverse relationship between type I-F CRISPR systems and *blaTEM-1* carriage is consistent with the suggestion that these bacterial immune systems play a role in shaping the antibiotic resistance profiles of *E. coli* populations.

### CRISPR array types as critical predictors of resistance: insights from penalised multivariate regression modelling

#### Predictors of trimethoprim resistance

Elastic net and adaptive lasso regression models were used to identify features associated with trimethoprim resistance. Both approaches achieved high predictive accuracy during model training, with the elastic net model achieving a cross-validated AUC of 0.93 and adaptive lasso achieving a similar AUC of 0.92, indicating excellent discriminatory ability. Elastic net retained 27 predictors, while adaptive lasso simplified the model to 16 significant features (Supplementary Table 7). Notably the presence of a CRISPR type I-F array was identified as a strong negative predictor of trimethoprim resistance (Fig. 7a), associated with a 99.995% decrease in the odds of resistance (OR=0.00005, β=-9.99) (Table 1). This aligns with its hypothesised role of limiting the acquisition of resistance genes through horizontal gene transfer. In contrast, other CRISPR array types were positive predictors of resistance. The odds of resistance increased nearly 10-fold for type I-E systems (OR=9.81, β=2.28), and over 15-fold for strains with an I-E system coupled with an orphan I-F array (OR=15.53, β=2.74).

**Figure 7.**
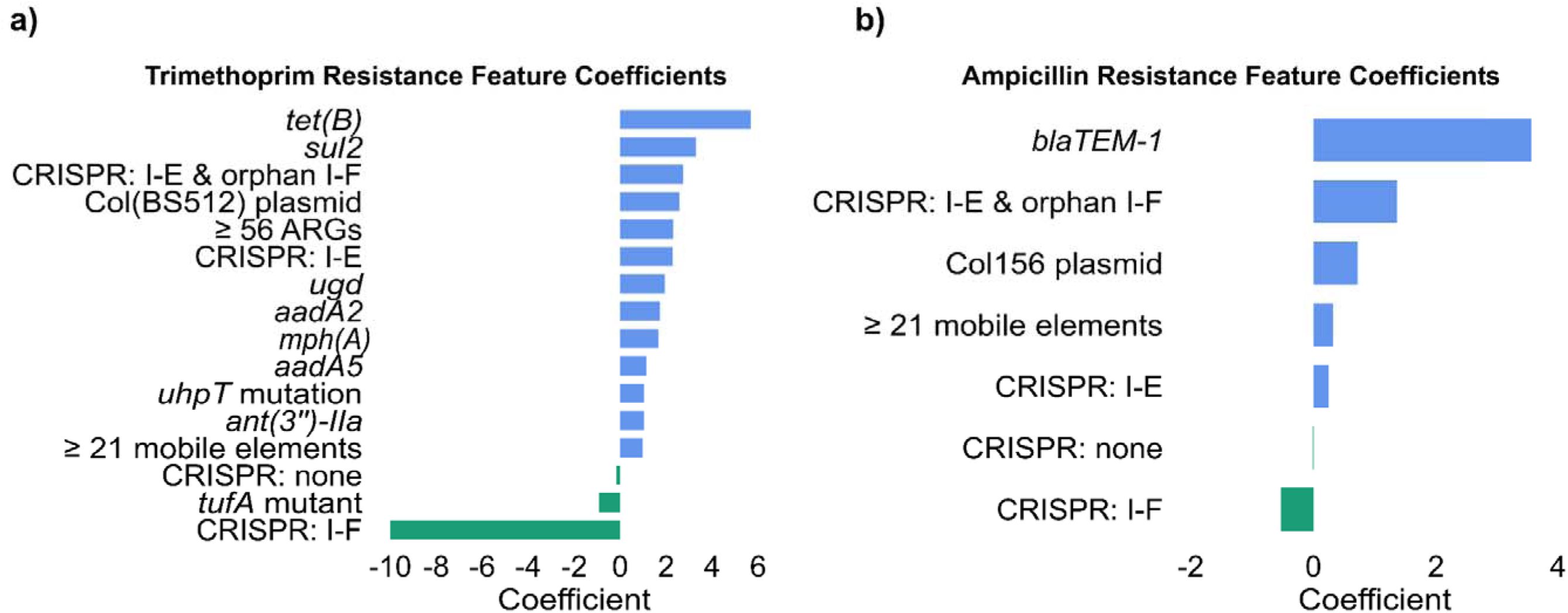
Predictors of antibiotic resistance identified by regularised regression in *E. coli* isolates. The plots display feature coefficients from elastic net models refined by adaptive lasso. Positive coefficients (blue) indicate a positive association with resistance, while negative coefficients (green) indicate a negative association. **a)** Trimethoprim resistance predictors: A CRISPR type I-F array was a strong negative predictor, while other CRISPR array types and resistance genes like *tet(B)*, *mph(A)*, and *ant(3”)-IIa* were positive predictors. **b)** Ampicillin resistance predictors: A CRISPR type I-F array was a negative predictor, while *blaTEM-1* was the strongest positive predictor.

**Table 1.**
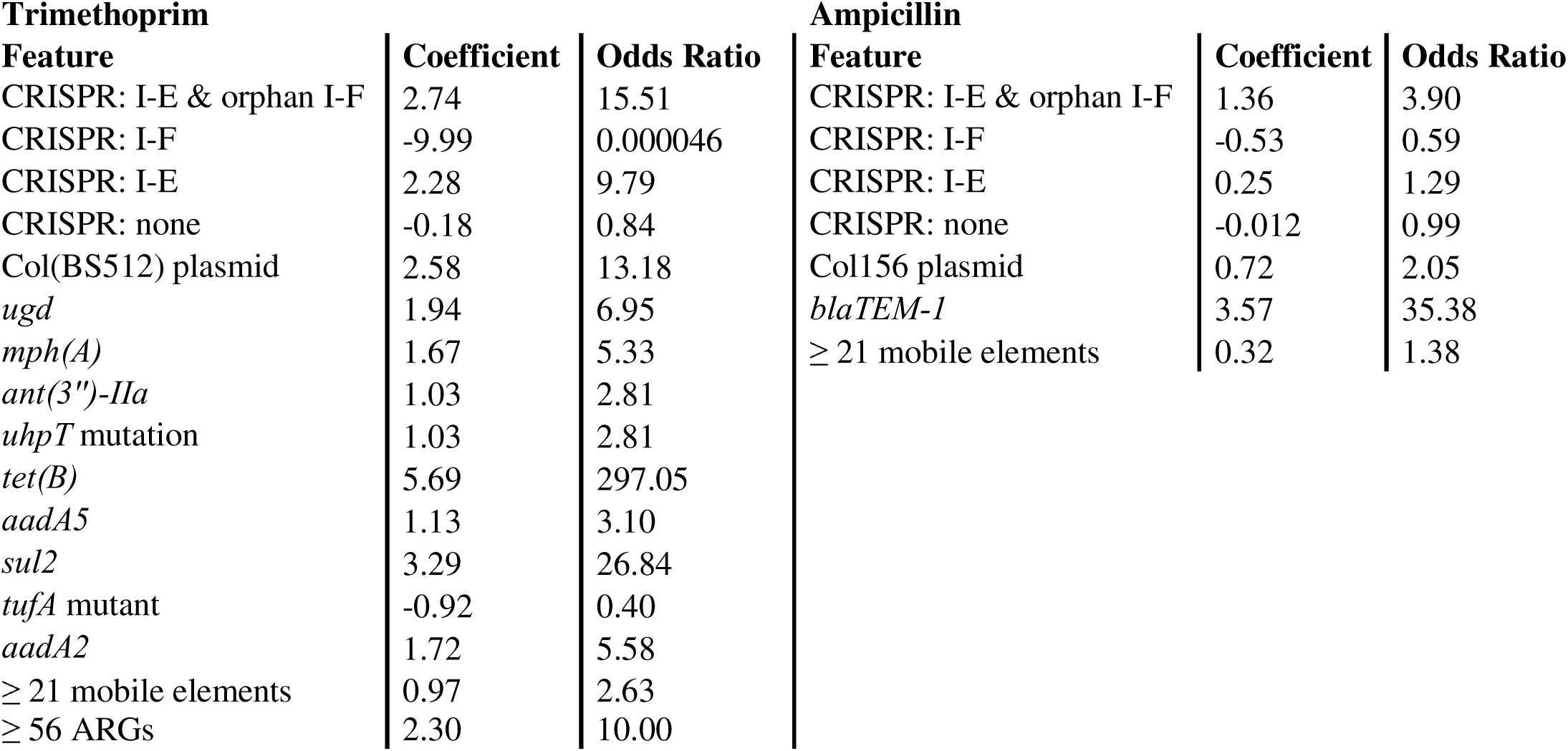
Coefficient Estimates and Odds Ratios for Predictors of Trimethoprim and Ampicillin Resistance in *E. coli* Isolates from regularised regressions.

| **Trimethoprim** |  |  | **Ampicillin** |  |  |
| --- | --- | --- | --- | --- | --- |
| **Feature** | **Coefficient** | **Odds Ratio** | **Feature** | **Coefficient** | **Odds Ratio** |
| CRISPR: I-E & orphan I-F | 2.74 | 15.51 | CRISPR: I-E & orphan I-F | 1.36 | 3.90 |
| CRISPR: I-F | -9.99 | 0.000046 | CRISPR: I-F | -0.53 | 0.59 |
| CRISPR: I-E | 2.28 | 9.79 | CRISPR: I-E | 0.25 | 1.29 |
| CRISPR: none | -0.18 | 0.84 | CRISPR: none | -0.012 | 0.99 |
| Col(BS512) plasmid | 2.58 | 13.18 | Col156 plasmid | 0.72 | 2.05 |
| *ugd* | 1.94 | 6.95 | *blaTEM-1* | 3.57 | 35.38 |
| *mph(A)* | 1.67 | 5.33 | ≥ 21 mobile elements | 0.32 | 1.38 |
| *ant(3'')-IIa* | 1.03 | 2.81 |  |  |  |
| *uhpT* mutation | 1.03 | 2.81 |  |  |  |
| *tet(B)* | 5.69 | 297.05 |  |  |  |
| *aadA5* | 1.13 | 3.10 |  |  |  |
| *sul2* | 3.29 | 26.84 |  |  |  |
| *tufA* mutant | -0.92 | 0.40 |  |  |  |
| *aadA2* | 1.72 | 5.58 |  |  |  |
| ≥ 21 mobile elements | 0.97 | 2.63 |  |  |  |
| ≥ 56 ARGs | 2.30 | 10.00 |  |  |  |

Key resistance-associated genes also significantly contributed to trimethoprim resistance. The presence of the gene *tet(B)* increased the odds of trimethoprim resistance 296-fold (β =3.29). Genes *mph(A)* and *ant(3”)-IIa* were also strong predictors of resistance, reflecting the genomic context of these genes within the composite transposons identified on the MDR plasmids of strains R52 and R60. The presence of *mph(A)* increased the odds of resistance by 430% (OR=5.3, β=1.67), while *ant(3”)-IIa* increased the odds by 180% (OR=2.8, β=1.03). Additionally, the presence of 56 or more resistance-related genes (OR=9.97, β=2.30), 21 or more mobile elements (OR=2.64, β=0.97), and the presence of a Col(BS512) plasmid (OR=1.57; β=0.45) were strongly associated with trimethoprim resistance.

The adaptive lasso model demonstrated strong classification performance on an independent global dataset (Table 2), with an AUC of 0.77 (Supplementary Fig. 3), confirming that the trends observed in the training data were consistent. The model was effective at identifying trimethoprim-resistant samples (sensitivity=72%) and sensitive strains (specificity=75%). This resulted in a balanced accuracy of 74% and an F1 score of 0.78. Full performance metrics are available in Supplementary Table 7.

**Table 2.**
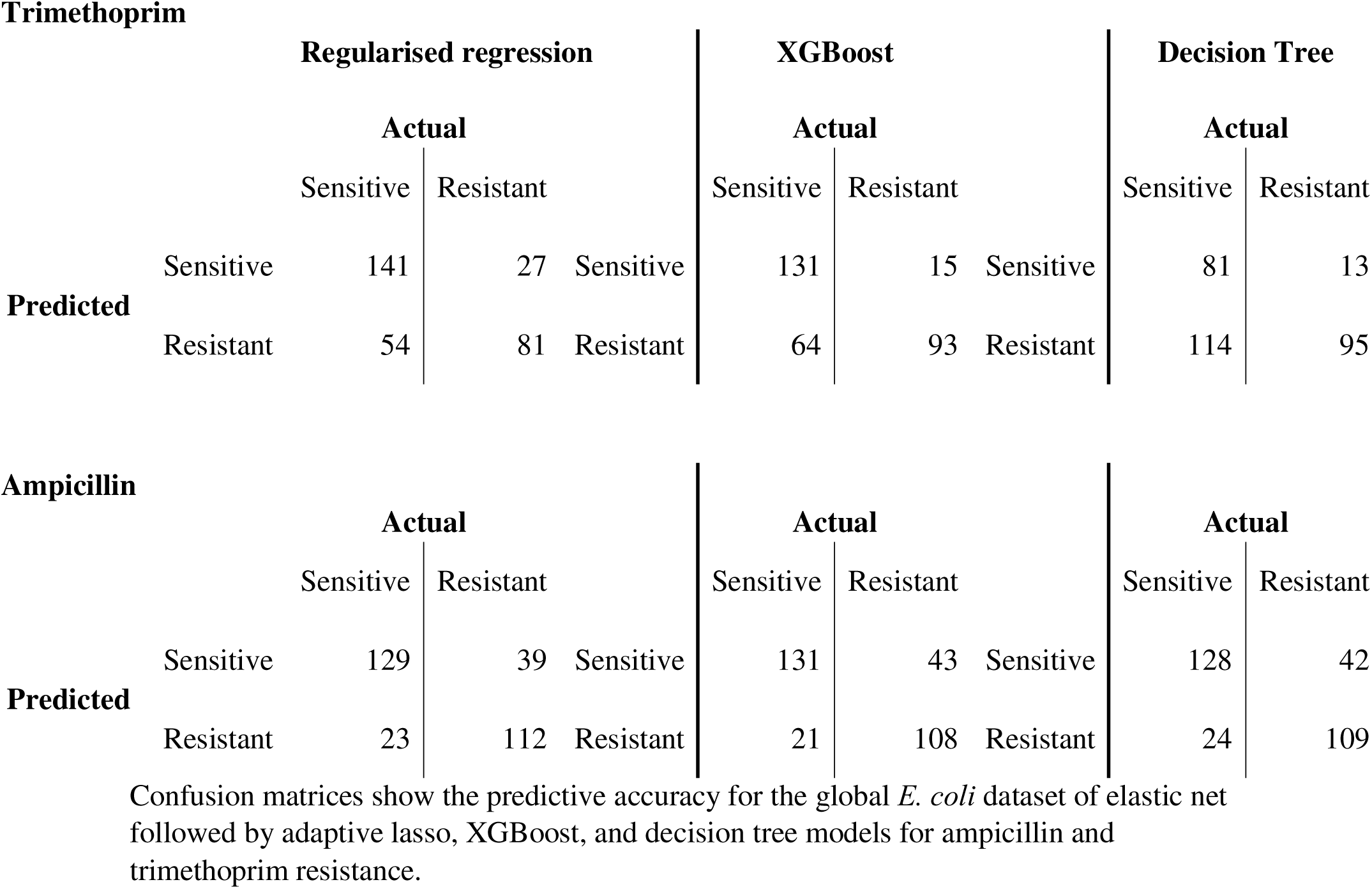
Performance of Machine Learning Models in Predicting Antibiotic Resistance.

#### Predictors of ampicillin resistance

Similarly, elastic net and adaptive lasso regression models were employed to identify predictors of ampicillin resistance. Both models demonstrated strong predictive accuracy during training, with the elastic net model yielding a cross-validated AUC of 0.86 and adaptive lasso achieving an AUC of 0.84. Elastic net retained 12 predictors, while adaptive lasso identified 7 significant features (Fig.7b; Supplementary Table 8). The presence of a CRISPR type I-F array again acted as a negative predictor, associated with a 41% decrease in the odds of ampicillin resistance (OR=0.59, β=-0.53) (Table 1). Conversely, CRISPR type I-E arrays had a slight positive association with resistance (OR=1.29, β=0.25), and this association was more pronounced when combined with an orphan I-F system (OR=3.89, β=1.36).

The *blaTEM-1* β-lactamase, an enzyme which inactivates ampicillin, identified on the R52 plasmid in this study, was the strongest predictor of ampicillin resistance (OR=35.56, β=3.56). The presence of 21 or more mobile elements increased the odds of resistance by 38% (OR=1.38, β=0.32), while the Col156 plasmid increased resistance odds by 105% (OR=2.05, β=0.72).

The adaptive lasso model demonstrated strong predictive accuracy with an AUC of 0.80 (Supplementary Fig. 4), confirming the robustness of the predictors identified in the training dataset (Supplementary Table 8). The best threshold, determined using Youden’s J, was found to be 0.74. In terms of classification performance, the model showed high sensitivity (84%), correctly identifying a large proportion of ampicillin-resistant strains, while maintaining a good specificity (74%) for sensitive strains. The model achieved a balanced accuracy of 80%, with an F1 score of 0.81, indicating a well-balanced performance across both classes.

Although the elastic net models provided slightly higher AUC values and retained a larger number of predictors, the adaptive lasso models demonstrated improved sparsity without compromising performance. Across both antibiotics, regularised regression modelling consistently highlighted the negative association of CRISPR type I-F arrays with resistance phenotypes, suggesting a conserved role in modulating resistance.

### XGBoost modelling: expanding the landscape of resistance genes and the persistent influence of CRISPR

Following the regularised regression analysis, we sought to validate these findings using an alternative machine learning approach. Specifically, we employed the XGBoost model due to its ability to capture complex, non-linear relationships within data. This allowed us to assess whether the key predictors identified by elastic net and adaptive lasso were robust to different modelling techniques and to explore additional factors potentially influencing resistance.

#### Predictors of trimethoprim resistance

The XGBoost model demonstrated strong discriminatory ability during model training, achieving a cross-validated AUC of 0.95. The model also demonstrated good generalisation performance on the independent global dataset (AUC = 0.81). While the model performed well when identifying trimethoprim-resistant samples (sensitivity=86%), identification of sensitive strains was somewhat reduced (specificity=67%). Although balanced accuracy remained high at 77%, the F1 score of 0.70 reflected the reduced specificity.

The XGBoost model identified several key predictors of trimethoprim resistance. The most influential factor was the presence of the *blaTEM-1* β-lactamase (Gain = 0.276). Several other resistance genes, including *ant(3”)-IIa* (Gain=0.116) and *sul2* (Gain=0.094), as well as the presence of 56 or more resistance-related genes (Gain=0.113), were strong positive predictors of resistance. While XGBoost modelling is able to explore potential non-linear relationships between genomic features and trimethoprim resistance, it can be more sensitive to class imbalance than regularised regression due to its focus on minimising error in the dominant class. Despite the relative rarity of the type I-F CRISPR array within the dataset, possession of this type was associated with decreased trimethoprim resistance (Gain = 0.006), supporting its hypothesised role in limiting the acquisition of resistance genes. Additionally, while not a strong predictor, the absence of any CRISPR array was also identified as a positive predictor (Gain = 0.011), suggesting that the overall CRISPR array status of the isolate may have an influence on trimethoprim resistance. The absence of any CRISPR array may contribute to trimethoprim resistance by leaving the bacterial isolate more vulnerable to the acquisition of resistance genes via horizontal gene transfer, as it lacks the adaptive immune defence provided by CRISPR-Cas systems ^19^.

The full list of XGBoost features and associated model gains are available in Supplementary Table 9.

#### Predictors of ampicillin resistance

For ampicillin resistance, XGBoost modelling identified far fewer predictors. Again, the most influential factor was the *blaTEM-1* β-lactamase (Gain=0.559). The possession of Col156 plasmids (Gain=0.130), *sul1* (Gain=0.115) and 21 or more mobile elements (Gain=0.094) were also positive predictors of ampicillin resistance.

This model performed consistently across training (AUC = 0.82) and independent validation (AUC = 0.81) datasets. The model demonstrated better performance in identifying ampicillin sensitive strains (specificity=86%) than ampicillin resistant strains (sensitivity=72%). This resulted in a balanced accuracy of 79% and an F1 score of 0.77.

The ampicillin resistance features and associated model gains identified by XGBoost are available in Supplementary Table 10.

### Decision tree modelling: simplicity and its limits in predicting resistance

Decision tree models offer an interpretable alternative to more complex machine learning approaches, allowing for a clear visualisation of potential factors driving resistance. However, their inherent simplicity can sometimes lead to oversimplification. Due to their susceptibility to picking up on collinear relationships, decision trees may prioritise predictive power over biological accuracy, potentially highlighting genes with limited direct involvement in the resistance mechanism.

#### Predictors of trimethoprim resistance

The decision tree model provided an interpretable alternative to more complex machine learning approaches, achieving a cross-validated training AUC of 0.83. When applied to the independent global validation dataset, the model retained reasonable discriminatory ability (AUC = 0.73), with high sensitivity (88%) but clearly reduced specificity (42%). This resulted in a balanced accuracy of 65% and an F1 score of 0.60, indicating that while the model was effective at identifying resistant isolates, it was less reliable in correctly identifying susceptible strains.

The model highlighted the importance of specific resistance genes, with the presence of *blaTEM-1* being the most influential factor. Isolates lacking *blaTEM-1* were further stratified by the presence of *ant(3”)-IIa* and *emrE*. Variable importance analysis also highlighted *mph(A)*, *sul1*, and *qacE*Δ*1* as relevant predictors, along with the presence of a type I-F CRISPR array, which was associated with reduced resistance probability. These findings align with those of the elastic net and XGBoost models, reinforcing their biological relevance. The complete tree structure and full variable importance rankings are provided in Supplementary Table 11 and Supplementary Fig. 5.

#### Ampicillin resistance: A simplified model

In contrast to trimethoprim resistance, the decision tree model for ampicillin resistance was overwhelmingly dominated by a single factor: the presence of the *blaTEM-1* gene. While the model maintained good performance on the training data (AUC = 0.80) and generalised similarly well to the independent validation dataset (AUC = 0.80), the resulting tree contained a single split on the *blaTEM-1* gene. This single node captured nearly all of the discriminatory signal, with the presence of *blaTEM-1* strongly predicting resistance, 96.7% (n=112/151), of *blaTEM-1*-positive isolates were resistant. The absence of *blaTEM-1* generally associated with antimicrobial susceptibility with only 15.2% (n=23/151) of ampicillin susceptible strains encoding the gene.

While the model had good performance, it highlights the potential for oversimplification inherent in this approach. Given the clear and consistent importance of *blaTEM-1*, as highlighted by the elastic net, adaptive lasso, and XGBoost models, the decision tree model offers limited additional insight into ampicillin resistance. The consistent and overwhelming importance of *blaTEM-1* suggests that diagnostic tests targeting this gene may be highly effective for predicting ampicillin resistance in this bacterial population. Full importance rankings are provided in Supplementary Table 11.

## Discussion

Our study provides compelling evidence that CRISPR array types are significant predictors of antibiotic resistance, particularly for resistance genes acquired through horizontal gene transfer. Using machine learning models, we demonstrate that type I-F arrays are strongly associated with susceptibility to antibiotics commonly encoded on mobile genetic elements, such as trimethoprim and ampicillin. Notably, CRISPR I-F arrays emerged as the strongest protective factor in regularised regression modelling, substantially reducing the odds of resistance. This finding supports the hypothesis I-F systems act as a barrier to the acquisition of ARGs via HGT. Conversely, type I-E arrays, especially in strains with additional orphan I-F arrays, correlate with increased resistance, suggesting a more complex or even potentially facilitative role in resistance acquisition for this CRISPR type. While counterintuitive, this may be best understood when viewed in light of the findings by Almendros et al ^21^, that orphan I-F arrays can prevent the uptake of complete, functional I-F CRISPR-Cas systems. The presence of orphan arrays could be an evolutionary strategy to increase genetic diversity through facilitating HGT and the acquisition of MGEs, particularly under the selective pressure of antibiotic exposure.

As expected, known ARGs, the presence of Col plasmids and a high burden of mobile genetic elements were strong contributors to resistance. Past studies have identified complex associations between CRISPR arrays, phylogroups, ARGs and MGEs within *E. coli*, but genetic interactions complicate the interpretability of these findings ^24,60,61^. Our multivariate modelling builds on these findings by accounting for these factors simultaneously, while leveraging the enhanced context provided by whole genome sequencing. This approach reduces potential confounding and offers a more integrated genomic perspective ^62^. Importantly, although our models were trained on bacterial strains collected from a single country in 2023, they performed with high predictive accuracy when applied to a global dataset, suggesting the associations may be generalisable. Altogether, these results point to a nuanced role of CRISPR systems in AMR, potentially driven by differences in the regulation of HGT.

Our finding that type I-F CRISPR arrays are associated with increased susceptibility to antibiotics commonly encoded on mobile genetic elements strongly supports the hypothesis that these systems act as a barrier to the acquisition of ARGs via HGT. This aligns with the reports that bacteria possessing CRISPR-Cas systems across a number of clinically relevant species, including *E. coli*, are less likely to harbor antibiotic resistance genes and MGEs compared to matched isolates ^17^, suggesting a protective role against the uptake of resistance determinants. While many prior studies drew upon large numbers of bacterial isolates when identifying this relationship, they have often been limited by their use of PCR-based methods and sequencing only of the CRISPR arrays ^24,58,61,63^, leaving the potential impact of the broader genomic context unexplored. These approaches lacked the resolution to fully clarify the interplay between different genetic elements and potential confounding factors.

In contrast, our genome-wide association study identifies CRISPR-Cas I-F systems as an independent predictor of antibiotic susceptibility, even after accounting for known ARGs, the presence of Col plasmids, mobile genetic elements, and phylogenetic relationships. This finding was further validated in a diverse external dataset, strengthening the robustness of our conclusions. These results highlight the potential protective role of CRISPR-Cas systems in controlling the spread of antibiotic resistance ^64^ and suggest that manipulating these systems could offer novel strategies for combating AMR ^65^. Furthermore, our study provides stronger statistical evidence for these associations, using a combination of genome-wide association and machine learning-based feature selection techniques.

Our findings highlight the complex relationship between CRISPR-Cas systems and antibiotic resistance, suggesting that CRISPR-Cas systems can act as both a barrier to and a facilitator of resistance acquisition, depending on the specific array type. Further research is warranted to fully elucidate the mechanisms underlying these associations. Specifically, a deeper comparison of the I-F and I-E systems is warranted, as they appear to exhibit contrasting roles in antibiotic resistance. Consistent with our modelling, CRISPR type I-E was more often associated with MDR strains in avian pathogenic *E. coli*, while I-F positive isolates have been linked to reduced resistance ^24,61^, although other studies have identified conflicting evidence. Research has also indicated that antibiotic resistance plasmids can still spread effectively within *E. coli* populations, even in the presence of CRISPR elements ^23^, and the impact of CRISPR-Cas systems on plasmid count is not consistent across bacterial species ^22^.

This discrepancy in the literature may reflect the divergent biological functions and regulatory states of CRISPR subtypes in *E. coli.* Type I-E systems are predominantly repressed and primarily target phage sequences ^60,66^, whereas I-F arrays are constitutively expressed and target both plasmids and phages ^20,60^, making them more likely to interfere with HGT.

Even within I-F systems, substantial heterogeneity exists. For example, CRISPR-associated transposons (CASTs) represent a specialised subtype, co-opted by Tn7-like transposons for site specific insertion ^67–69^. These CASTs typically lack key components of canonical I-F systems, including the Cas1 and Cas2 proteins required for spacer acquisition and the Cas3 nuclease responsible for target site cleavage ^68,69^. As a result, they likely do not have the functional capacity to prevent resistance gene acquisition. The presence of orphan or CAST-type I-F arrays may therefore account for the variability observed across studies. Moreover, prior studies relying on PCR-based screening may have inadvertently misclassified these subtypes, contributing to inconsistent findings. Moving forward, clarifying the expression, activity, and functional diversity of CRISPR subtypes will be essential to understanding their mechanistic role in AMR dynamics.

These findings have notable translational implications. The robust and subtype-specific association between CRISPR-Cas systems and antibiotic susceptibility suggests that CRISPR array typing may serve as a novel genomic biomarker for AMR risk stratification. Unlike conventional resistance gene screening, CRISPR profiling provides a broader lens on the bacterial strain’s potential to resist horizontal gene acquisition. CRISPR typing may add predictive value to genome-based antimicrobial susceptibility testing (g-AST) workflows ^70^, particularly as sequencing becomes increasingly integrated into clinical microbiology ^71^. Identifying strains with protective CRISPR types, such as I-F, may also support epidemiological surveillance and risk prediction efforts.

Beyond diagnostics, this work lays a conceptual foundation for future therapeutic strategies that harness or engineer CRISPR-Cas activity to limit the spread of resistance plasmids. Such strategies may complement antimicrobial stewardship by targeting resistance propagation at the genomic level. Further characterisation and leveraging of CRISPR-Cas activity could contribute to the development of precision antimicrobial tools that limit resistance gene dissemination ^65^. As CRISPR-based technologies advance toward clinical utility, understanding subtype-specific activity will be essential for both safe implementation and targeted microbial control.

## Conclusions

Our study provides strong evidence that CRISPR-Cas systems play a functional role in shaping antibiotic resistance profiles in *E. coli*. Specifically, we demonstrate that type I-F CRISPR arrays are an independent and consistent predictors of susceptibility to antibiotics commonly acquired through horizontal gene transfer, including trimethoprim and ampicillin. Conversely, type I-E arrays, particularly when co-occurring with orphan I-F arrays, are associated with increased resistance. These associations were validated in a globally diverse dataset, reinforcing their generalisability.

Together, these findings highlight the dualistic nature of CRISPR-Cas systems in modulating AMR dynamics and point toward new opportunities for integrating CRISPR profiling into genomic surveillance, predictive, diagnostics, or even precision antimicrobial interventions. Future work should focus on resolving the mechanistic pathways linking CRISPR subtype to mobile element dynamics, and evaluating the feasibility of CRISPR-guided interventions to contain resistance gene dissemination in clinical settings.

## Data availability

The sequencing data generated during the current study were deposited in NCBI under Bioproject PRJNA1279687. All other WGS were obtained from the SRA public repository and accession numbers are available in the Supplementary Table 3.

## Author contributions

A.M.Y. conducted all analyses, wrote the manuscript and performed bacterial DNA preparation. P.H. provided critical guidance in statistical analysis and verified the methods used in the manuscript. F.L. guided and prepared bacterial DNA for genome sequencing. M.C.W. organized and provided bacterial isolates and antibiotics susceptibility data for all Australian *E. coli* isolates in this manuscript. A.T. conducted bacterial genome sequencing S.M.R.: conceived the project together with L.Z. who also supervised the project. All authors provided critical feedback to contributed to manuscript editing.

## Acknowledgements

This project was supported by research fundings awarded to Associate Professor Li Zhang by University of New South Wales (Grant no. PS46772).

