## Supplementary Figures for "A Machine Learning Approach Reveals CRISPR-Cas I-F as a Genomic Marker of Antibiotic Susceptibility in Uropathogenic *E. coli*"

**Supplementary Table 2 Sequencing and assembly metrics statistics for long-read hybrid assemblies of *E. coli* isolates R52 and R60.**

|  | **Sequencing** | | | | | **Assembly** | | | | | |
| --- | --- | --- | --- | --- | --- | --- | --- | --- | --- | --- | --- |
| **ID** | **# reads (×10³)** | **Total bases sequenced (Gbp)** | **Read N50 (kb)** | **Mean read length (kb)** | **Mean Q** | **# contigs ≥1 kb** | **Total length (Mb)** | **Largest contig (Mb)** | **GC (%)** | **N50 (kb)** | **L50** |
| R52 | 153.6 | 1.77 | 27.9 | 11.5 | 12.3 | 2 | 5.33 | 5.21 | 50.45 | 5208 | 1 |
| R60 | 140.5 | 1.29 | 20 | 9.21 | 13.2 | 5 | 5.32 | 5.17 | 50.79 | 5167 | 1 |

This table presents genome assembly metrics for two *E. coli* isolates (R52 and R60) sequenced using Oxford Nanopore long-read technology and polished with Illumina short reads. Hybrid assemblies were generated using Trycycler, with polishing performed via Medaka and Polypolish. Raw-read metrics were computed with NanoStat v1.6.0 and include the number of passed reads, bases sequenced, N50, mean read length and mean Q. Assembly statistics were computed using QUAST v5.2.0 and include the number of contigs ≥1000 bp, total assembly length, largest contig, GC content, N50, and L50. Both assemblies reflect high contiguity and near-complete genome resolution.
Definitions: N50 = length of the contig at which 50% of the total assembly length is reached; Mean Q = mean per-base Phred quality score across all reads L50 = number of contigs contributing to N50.

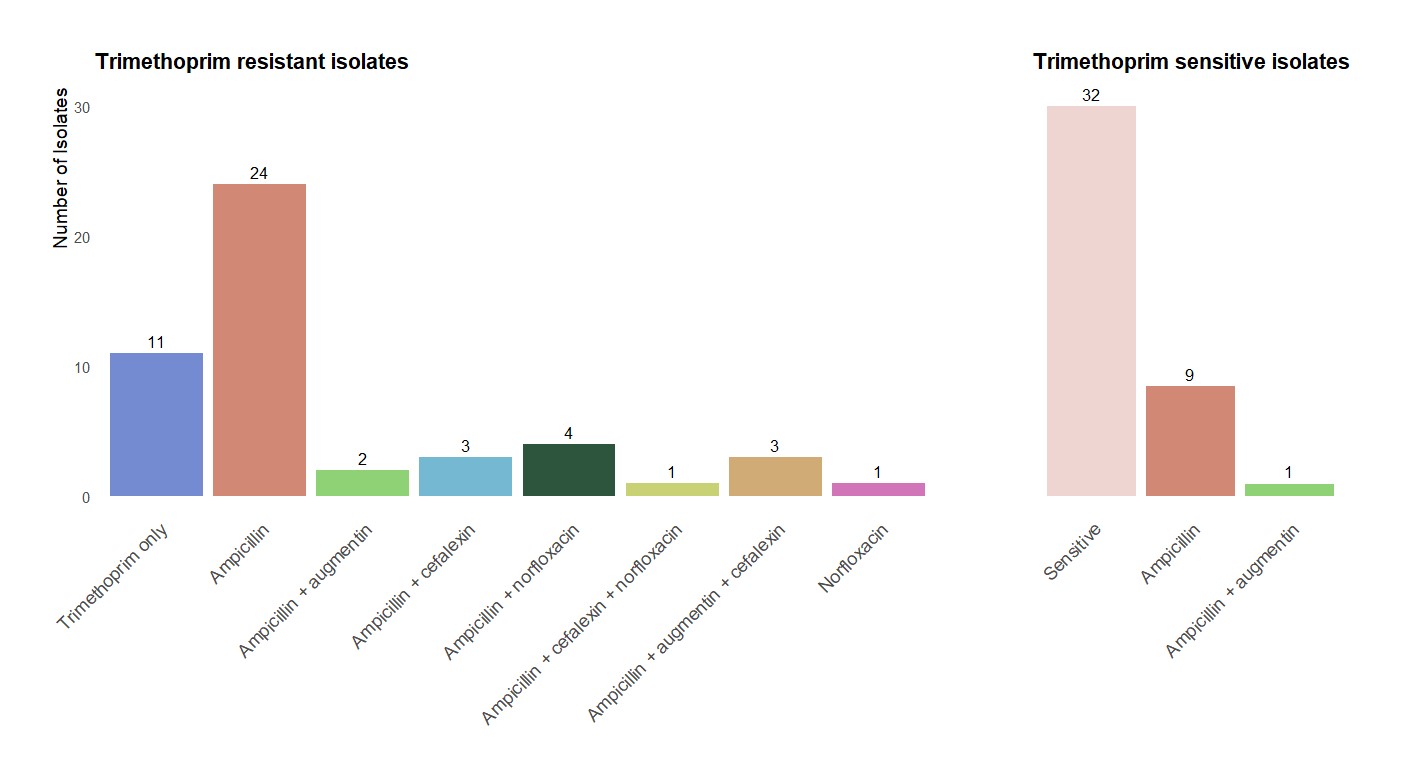
**Supplementary Figure 1 Antimicrobial resistance profiles and co-resistance patterns of *E. coli* sequence isolates**. A bar graph depicting the number of isolates resistant to ampicillin, amoxicillin with clavulanate, cefalexin, and norfloxacin. A high degree of co-resistance was observed between trimethoprim and ampicillin, with approximately 75% of trimethoprim-resistant isolates also resistant to ampicillin. There were 32 isolates were susceptible to all antibiotics tested, and no isolates displayed resistance to nitrofurantoin

**Supplementary Table 5 Hyperparameter grid for XGBoost model tuning.**

| **Hyperparameter** | **Values Tested** | **Description** |
| --- | --- | --- |
| Number of Boosting Iterations | 50, 100 | Total number of trees (iterations) used to fit the model. |
| Maximum Tree Depth | 4, 6, 8 | Maximum depth of each tree; controls model complexity. |
| Learning Rate | 0.01, 0.1, 0.3 | Step size shrinkage used to prevent overfitting. |
| Minimum Loss Reduction for Tree Split | 0, 1 | Minimum reduction in loss required to make a further split in a decision tree node. |
| Column Subsampling Ratio per Tree | 0.7, 1 | Fraction of predictor variables (features) randomly sampled for each tree. |
| Minimum Sum of Instance Weight per Child Node | 1, 5 | Minimum sum of instance weight (Hessian) in a child node; higher values make the algorithm more conservative. |
| Row Subsampling Ratio per Boosting Round | 0.7, 1 | Fraction of observations randomly sampled for each boosting round; reduces overfitting. |

This table outlines the hyperparameters and corresponding values tested during model optimization using a grid search. The search was conducted using the expand.grid() function in R, and model performance was assessed using 8-fold cross-validation.

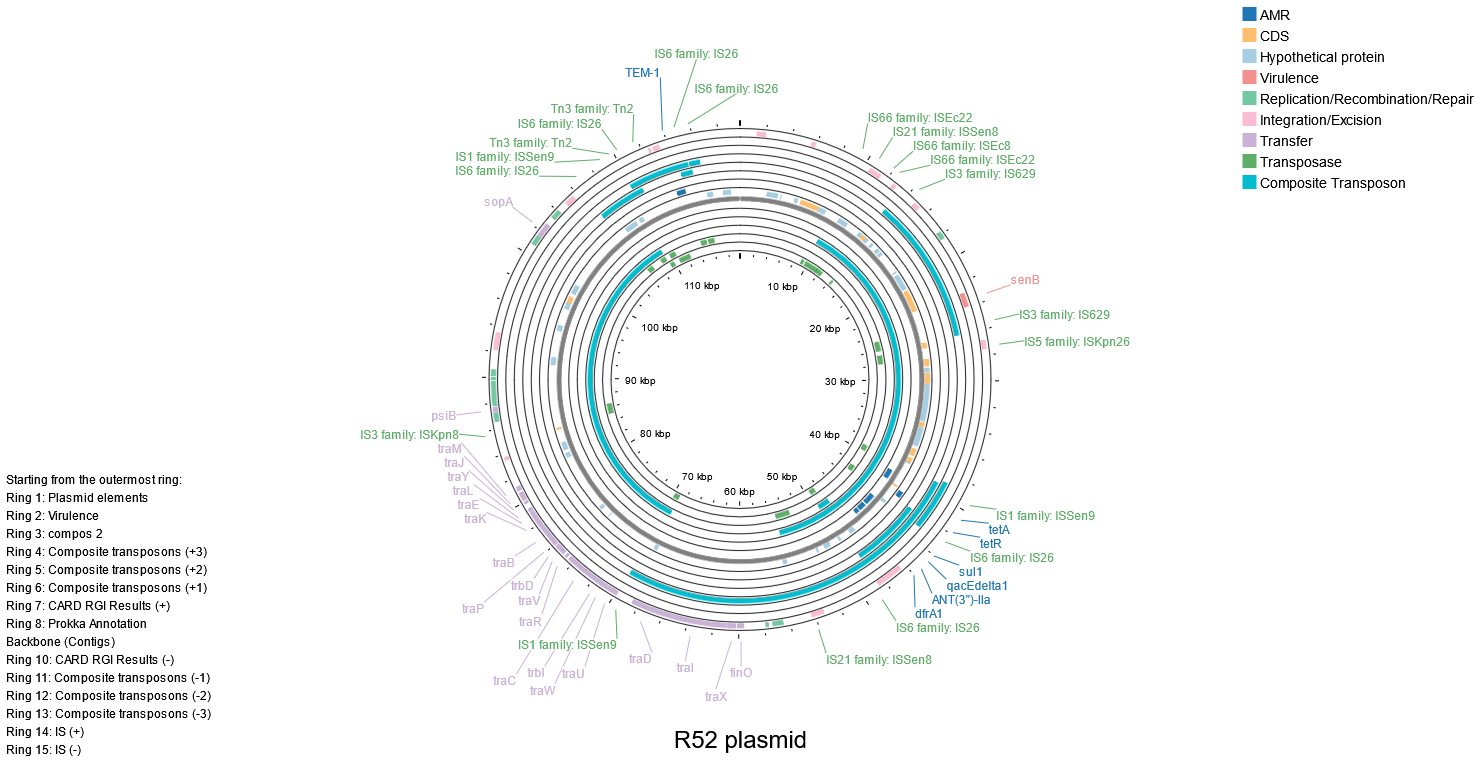

**b)**

**a)**

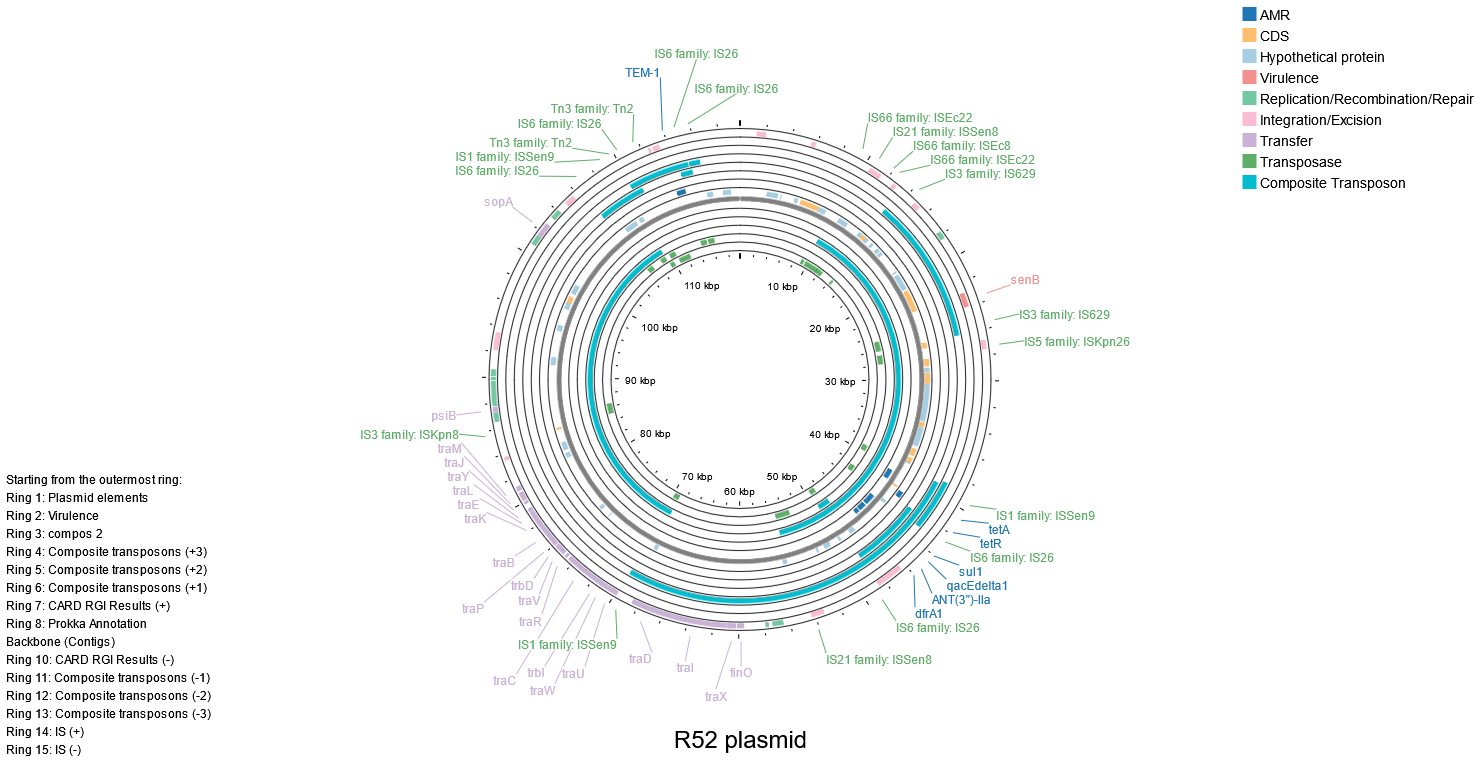

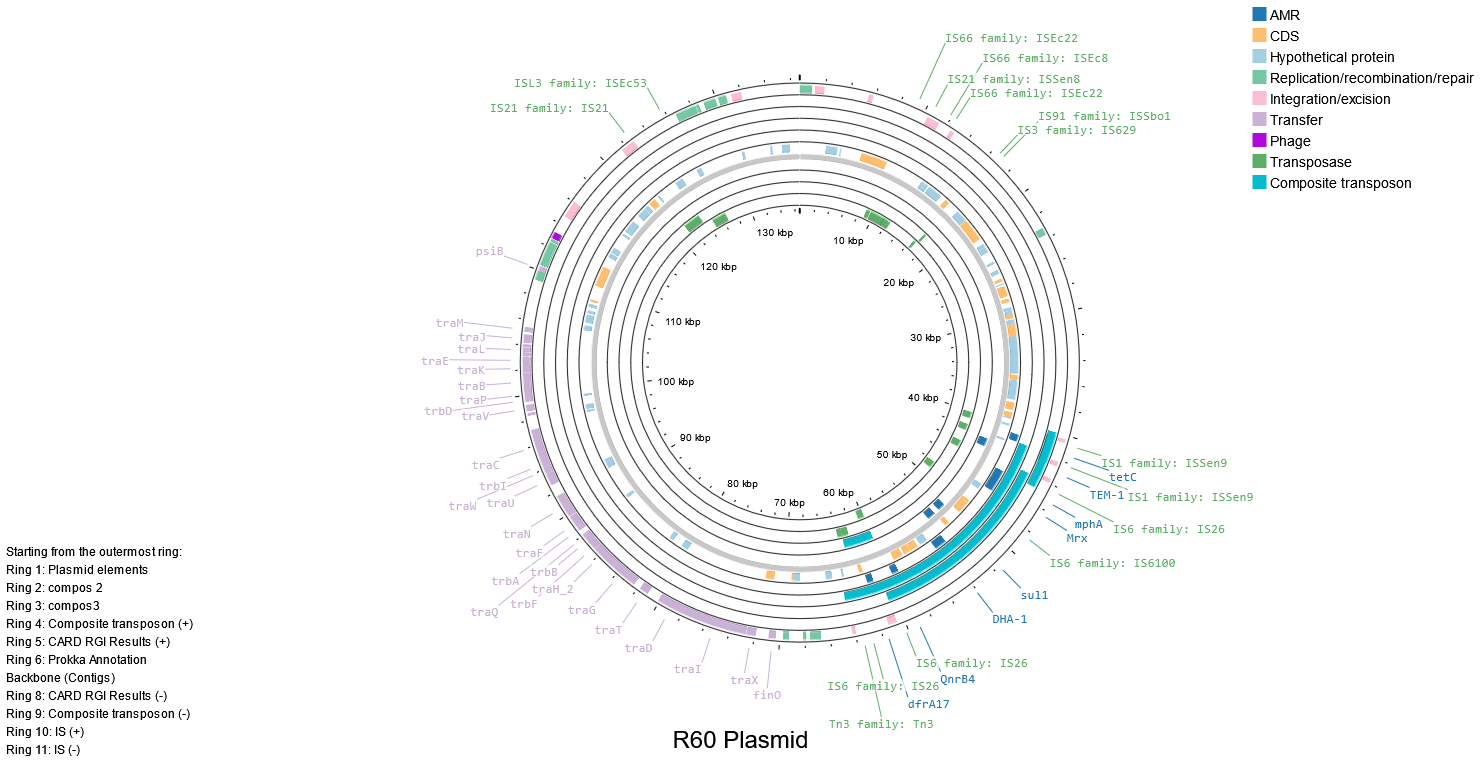

**Supplementary Figure 2 Annotated multi-drug resistant plasmids containing composite transposons in R52 and R60 Isolates. A)** *E. coli* R52 119kb FIA-type **B)** R60 FII-type 136kb plasmid with hypermobile resistance region. Colour-coded features represent resistance genes, insertion sequences, transposases, toxin–antitoxin systems, and other mobile elements as indicated in the figure legend. Plasmid similarity was confirmed using BLASTn against publicly available sequences (see main text).

**Supplementary Table 7 Trimethoprim resistance model features and performance metrics from elastic net regression.**

**Panel A** Features selected by elastic net model for trimethoprim resistance prediction

| **Feature** | **Coefficient** | **Odds Ratio** |
| --- | --- | --- |
| CRISPR: I-E & orphan I-F | 1.79 | 5.97 |
| CRISPR: I-F | -10.12 | 0.000040 |
| CRISPR: I-E | 1.50 | 4.50 |
| CRISPR: none | -0.10 | 0.90 |
| Phylogroup: D | 0.02 | 1.03 |
| Col(BS512) plasmid | 1.32 | 3.74 |
| *blaEC-8* | 0.41 | 1.51 |
| *ugd* | 1.48 | 4.38 |
| *mphA* | 0.96 | 2.60 |
| *cyaA* | 0.32 | 1.38 |
| *emrE* | -0.10 | 0.90 |
| *ant(3'')-IIa* | 1.09 | 2.96 |
| PtsI | 0.39 | 1.48 |
| *uhpT* mutation | 0.73 | 2.08 |
| *tet(B)* | 2.72 | 15.15 |
| *aadA5* | 1.05 | 2.87 |
| *tet(A)* | 0.57 | 1.78 |
| *aph(3'')-Ib* | 0.83 | 2.29 |
| *sul2* | 1.16 | 3.18 |
| *aac(3)-IId* | 0.42 | 1.52 |
| *gyrA* | -0.12 | 0.89 |
| *parC* | 0.29 | 1.34 |
| *tufA* mutant | -0.82 | 0.44 |
| *aph6)-Id* | 0.64 | 1.89 |
| *aadA2* | 0.94 | 2.56 |
| ≥ 21 mobile elements | 0.81 | 2.26 |
| ≥ 56 antibiotic resistance genes | 1.28 | 3.58 |

**Panel B** Predictive performance of the elastic net regression model on the global validation dataset

| **Metric** | **Value** |
| --- | --- |
| Accuracy | 0.73 |
| Sensitivity | 0.73 |
| Specificity | 0.75 |
| Positive predictive value | 0.84 |
| Negative predictive value | 0.60 |
| Balanced accuracy | 0.74 |

This table presents the final model output from adaptive lasso regression predicting trimethoprim resistance in *E. coli* isolates. Panel A lists the genomic features retained after regularisation, including CRISPR array types, resistance genes, and mobile element counts, along with their regression coefficients and odds ratios. Panel B summarizes the model’s predictive performance on the independent global validation dataset, including accuracy, sensitivity, specificity, and balanced accuracy.
Cut-offs used for mobile elements (≥21) and resistance genes (≥56) were defined based on the distribution in the training data.

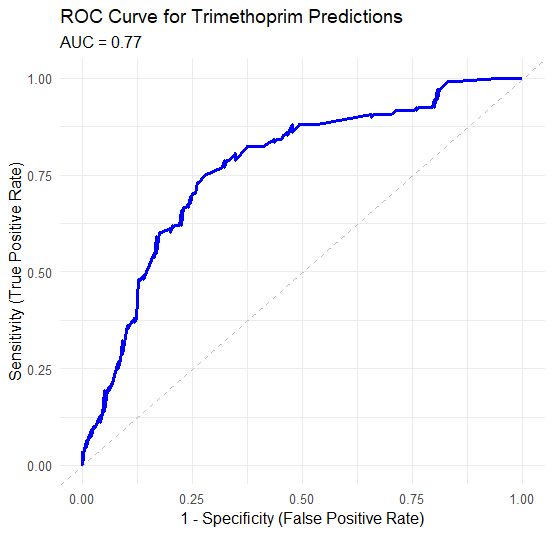

**Supplementary Figure 3 ROC Curve for Adaptive Lasso Model Predicting Trimethoprim Resistance in *E. coli*.** Receiver operating characteristic (ROC) curve showing the performance of the adaptive lasso logistic regression model for predicting trimethoprim resistance in *Escherichia coli*. The model was trained on isolates from the local dataset and validated on an independent, globally sourced cohort. The area under the ROC curve (AUC) was 0.77, indicating strong discriminatory ability. The optimal classification threshold was determined using Youden’s J statistic at 0.52.

**Supplementary Table 8 Ampicillin resistance model features and performance metrics from elastic net regression.**

**Panel A** Features selected by elastic net model for ampicillin resistance prediction

| **Feature** | **Coefficient** | **Odds Ratio** |
| --- | --- | --- |
| CRISPR: I-E & orphan I-F | 1.16 | 3.19 |
| CRISPR: I-F | -0.95 | 0.39 |
| CRISPR: I-E | 0.31 | 1.37 |
| CRISPR: none | -0.01 | 0.99 |
| Col156 plasmid | 0.45 | 1.57 |
| *ugd* | 0.25 | 1.29 |
| *mphA* | 0.43 | 1.54 |
| *sul1* | 0.21 | 1.23 |
| *blaTEM-1* | 1.44 | 4.24 |
| *qacEΔ1* | 0.14 | 1.15 |
| *tet(A)* | 0.14 | 1.15 |
| ≥ 21 mobile elements | 0.35 | 1.43 |

**Panel B** Predictive performance of the elastic net regression model on the global validation dataset

| **Metric** | **Value** |
| --- | --- |
| Accuracy | 0.795 |
| Sensitivity | 0.849 |
| Specificity | 0.742 |
| Positive predictive value | 0.768 |
| Negative predictive value | 0.8296 |
| Balanced accuracy | 0.795 |

This table presents the final model output from adaptive lasso regression predicting ampicillin resistance in *E. coli* isolates. Panel A lists the genomic features retained after regularisation, including CRISPR array types, resistance genes, plasmid presence and mobile element counts, along with their regression coefficients and odds ratios. Panel B summarizes the model’s predictive performance on the independent global validation dataset, including accuracy, sensitivity, specificity, and balanced accuracy.
The cut-off used for mobile elements (≥21) was defined based on the distribution in the training data.

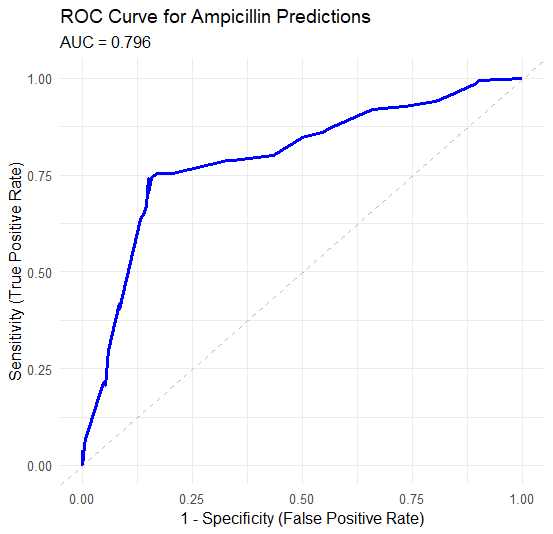

**Supplementary Figure 4 ROC Curve for Adaptive Lasso Model Predicting Ampicillin Resistance in *E. coli*.** Receiver operating characteristic (ROC) curve showing the performance of the adaptive lasso logistic regression model for predicting ampicillin resistance in *Escherichia coli*. The model was trained on isolates from the local dataset and validated on an independent, globally sourced cohort. The area under the ROC curve (AUC) was 0.796, indicating strong discriminatory ability. The optimal classification threshold was determined using Youden’s J statistic at 0.74.

**Supplementary Table 9 XGBoost Feature Importance and Performance Metrics for Predicting Trimethoprim Resistance.**

**Panel A:** Feature Importance

| **Rank** | **Feature** | **Gain** | **Cover** | **Frequency** |
| --- | --- | --- | --- | --- |
| 1 | *blaTEM-1* | 0.2760 | 0.1283 | 0.10 |
| 2 | *ant(3'')-IIa* | 0.1160 | 0.0780 | 0.10 |
| 3 | ≥ 56 antibiotic resistance genes | 0.1131 | 0.1270 | 0.10 |
| 4 | *sul2* | 0.0945 | 0.1383 | 0.10 |
| 5 | *blaEC-5* | 0.0772 | 0.0845 | 0.10 |
| 6 | *tet(B)* | 0.0627 | 0.1075 | 0.08 |
| 7 | *cyaA* mutation | 0.0388 | 0.0229 | 0.02 |
| 8 | ≥21 mobile elements | 0.0351 | 0.0365 | 0.06 |
| 9 | B2 phylogroup | 0.0291 | 0.0334 | 0.04 |
| 10 | IncFII plasmid | 0.0279 | 0.0240 | 0.04 |
| 11 | *emrE* | 0.0264 | 0.0463 | 0.04 |
| 12 | *mphA* | 0.0260 | 0.0510 | 0.04 |
| 13 | IncFIA plasmid | 0.0169 | 0.0053 | 0.02 |
| 14 | *mdtM* | 0.0166 | 0.0183 | 0.04 |
| 15 | CRISPR: none | 0.0111 | 0.0067 | 0.02 |
| 16 | A phylogroup | 0.0082 | 0.0182 | 0.02 |
| 17 | *ptsI* mutation | 0.0069 | 0.0204 | 0.02 |
| 18 | CRISPR: I-F | 0.0064 | 0.0225 | 0.02 |
| 19 | Col156 plasmid | 0.0063 | 0.0174 | 0.02 |
| 20 | *gyrA* mutation | 0.0049 | 0.0135 | 0.02 |

**Panel B:** Model Performance on Global Validation Set

| **Metric** | **Value** |
| --- | --- |
| Accuracy | 0.739 |
| Sensitivity | 0.861 |
| Specificity | 0.672 |
| Positive predictive value | 0.592 |
| Negative predictive value | 0.897 |
| Balanced accuracy | 0.767 |

This table summarizes the most influential genomic features and validation performance of the XGBoost classifier trained to predict trimethoprim resistance in *E. coli*. Feature importance was ranked based on Gain (contribution to model accuracy), with Cover and Frequency also reported. Model performance was evaluated on an independent global dataset. The optimal threshold was defined using Youden’s J statistic at 0.548.

**Supplementary Table 10 XGBoost Feature Importance and Validation Performance for Predicting Ampicillin Resistance in *E. coli*.**

**Panel A:** Feature Importance

| **Rank** | **Feature** | **Gain** | **Cover** | **Frequency** |
| --- | --- | --- | --- | --- |
| 1 | *blaTEM-1* | 0.5591 | 0.2498 | 0.2045 |
| 2 | Col156 plasmid | 0.1300 | 0.2393 | 0.2273 |
| 3 | *sul1* | 0.1146 | 0.0598 | 0.0455 |
| 4 | ≥21 mobile elements | 0.0935 | 0.2460 | 0.2727 |
| 5 | *qacEΔ1* | 0.0463 | 0.0299 | 0.0227 |
| 6 | *blaEC-5* | 0.0311 | 0.0815 | 0.1136 |
| 7 | *mdtM* | 0.0133 | 0.0542 | 0.0682 |

**Panel B:** Model Performance on Global Validation Set

| **Metric** | **Value** |
| --- | --- |
| Accuracy | 0.789 |
| Sensitivity | 0.715 |
| Specificity | 0.862 |
| Positive predictive value | 0.837 |
| Negative predictive value | 0.753 |
| Balanced accuracy | 0.789 |

This table presents the top genomic features and validation performance of the XGBoost classifier trained to predict ampicillin resistance in *Escherichia coli*. Feature importance was calculated based on Gain (contribution to accuracy), with Cover and Frequency also shown.
The model was trained on the local dataset and validated using an independent global cohort. The classification threshold was selected using Youden’s J statistic at 0.663.

**Supplementary Table 11 Decision tree variable importance rankings for predicting trimethoprim and ampicillin resistance.**

|  | **Trimethoprim** | | **Ampicillin** | |
| --- | --- | --- | --- | --- |
| **Rank** | **Variable** | **Importance** | **Variable** | **Importance** |
| 1 | ≥ 56 antibiotic resistance genes | 100.00 | *blaTEM-1* | 18.14 |
| 2 | *blaTEM-1* | 89.24 | *sul1* | 10.28 |
| 3 | *mph(A)* | 70.82 | *qacEΔ1* | 9.67 |
| 4 | *sul1* | 62.46 | *mph(A)* | 9.07 |
| 5 | *qacEΔ1* | 56.79 | *tet(A)* | 7.26 |
| 6 | Col156 | 39.18 | *aac(3)-IId* | 5.44 |
| 7 | *ant(3'')-IIa* | 33.74 |  |  |
| 8 | *emrE* | 26.44 |  |  |
| 9 | *cyaA* mutation | 26.44 |  |  |
| 10 | Phylogroup D | 24.75 |  |  |
| 11 | *blaEC-8* | 24.75 |  |  |
| 12 | CRISPR: I-F | 20.81 |  |  |
| 13 | ≥21 mobile elements | 17.95 |  |  |

This table reports the top-ranked predictors for antimicrobial resistance as identified by decision tree models trained on the Australian *E. coli* dataset. Variables are ranked by their relative importance (scaled or raw, depending on the model), with higher values indicating greater contribution to resistance prediction. While the ampicillin model was dominated by a single predictive feature (*blaTEM-1*), multiple features contributed to trimethoprim resistance predictions. Note that decision trees are inherently unstable and rankings may vary slightly across model iterations.

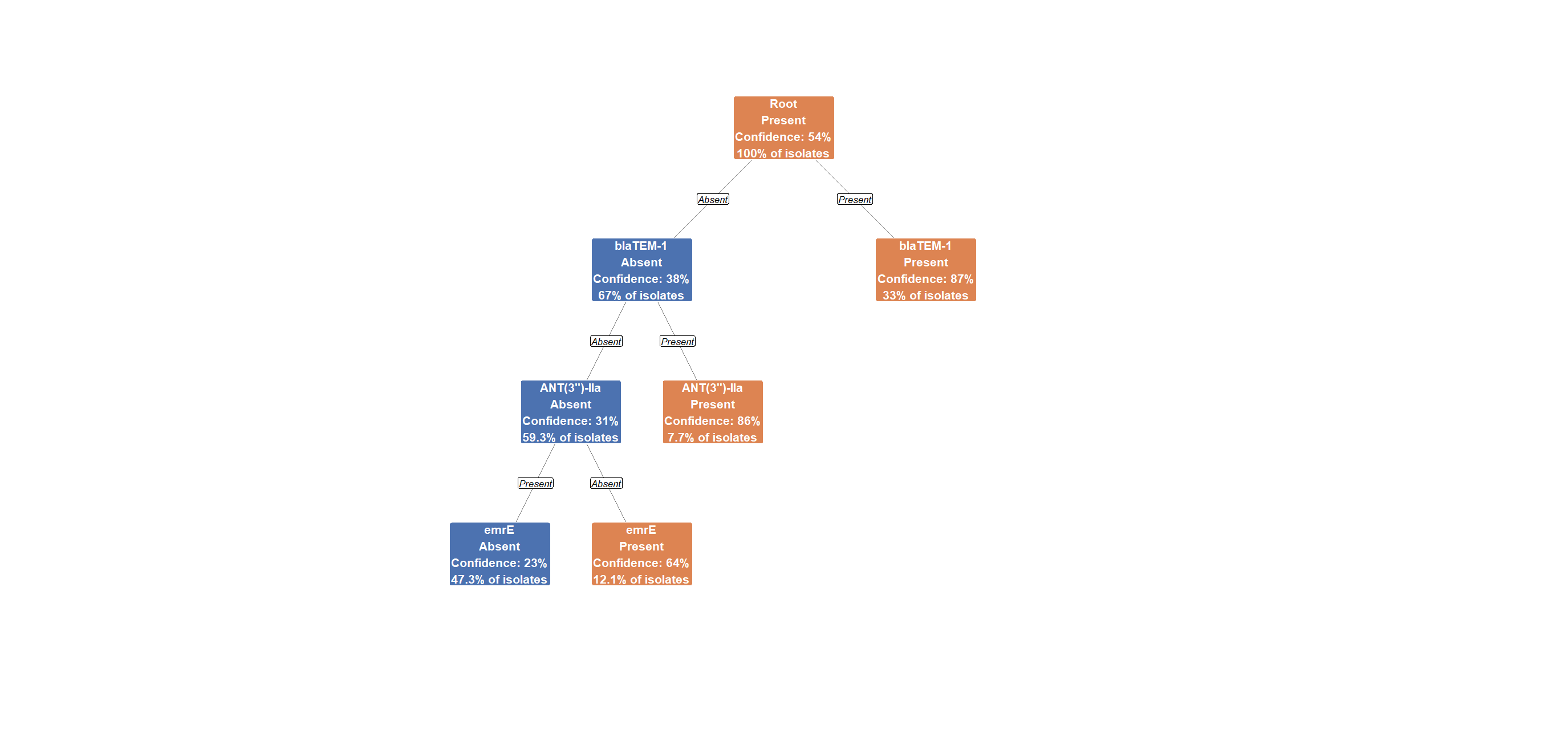

**Supplementary Figure 5** **Decision tree model predicting trimethoprim resistance in uropathogenic *E. coli***. This figure shows the structure of the decision tree trained on the Australian *E. coli* dataset to classify trimethoprim resistance based on genomic features. Nodes represent binary decision points using the presence or absence of key antimicrobial resistance genes and other genomic elements. Each node displays the predicted class (Present or Absent), the confidence level of the prediction, and the proportion of isolates within that branch. The root split occurs on *blaTEM-1*, the most influential feature in the model. Subsequent splits include *ant(3'')-IIa and emrE*, contributing to further classification. Orange nodes indicate a predicted resistant phenotype, and blue nodes indicate susceptibility. While the model offers visual interpretability, its reduced specificity relative to more complex models (e.g., XGBoost) suggests limitations in capturing the full complexity of resistance phenotypes.
